# Multi-tissue landscape of somatic mtDNA mutations indicates tissue specific accumulation and removal in aging

**DOI:** 10.1101/2022.08.30.505884

**Authors:** Monica Sanchez-Contreras, Mariya T. Sweetwyne, Kristine A. Tsantilas, Jeremy A. Whitson, Matthew D. Campbell, Brendan F. Kohrn, Hyeon Jeong Kim, Micheal J. Hipp, Jeanne Fredrickson, Megan M. Nguyen, James B. Hurley, David J. Marcinek, Peter S. Rabinovitch, Scott R. Kennedy

**Author notes:** authors contributed equally (Monica Sanchez-Contreras, Mariya T. Sweetwyne).

## Abstract

Accumulation of somatic mutations in the mitochondrial genome (mtDNA) during aging has long been proposed as a possible mechanism of mitochondrial and tissue dysfunction. A thorough characterization of age-associated mtDNA somatic mutations has been hampered by the limited ability to detect low frequency mutations. Here, we used Duplex Sequencing on eight tissues of an aged mouse cohort to detect >89,000 independent somatic mtDNA mutations and show significant tissue-specific increases during aging across all tissues examined which did not correlate with mitochondrial content and tissue function. G→A/C→T substitutions, indicative of replication errors and/or cytidine deamination, were the predominant mutation type across all tissues and increased with age, whereas G→T/C→A substitutions, indicative of oxidative damage, were the second most common mutation type, but did not increase with age regardless of tissue. We also show that clonal expansions of mtDNA mutations with age is tissue and mutation type dependent. Unexpectedly, mutations associated with oxidative damage rarely formed clones in any tissue and were significantly reduced in the hearts and kidneys of aged mice treated at late age with Elamipretide or nicotinamide mononucleotide. Thus, the lack of accumulation of oxidative damage-linked mutations with age indicates a life-long dynamic clearance of either the oxidative lesions or mtDNA genomes harboring oxidative damage.

## INTRODUCTION

Genetic instability is a hallmark of aging (López-Otín et al., 2013). A mechanistic link between somatic mutations and age-related diseases such as cancer is clear, but their importance in other aging phenotypes, long hypothesized, is poorly understood (Zhang and Vijg, 2018). Recent surveys of non-diseased somatic tissues have shown that mutations are pervasive in the nuclear genome (nDNA), increase with age, and vary considerably between tissues (Abascal et al., 2021; Li et al., 2021). Additionally, these nDNA mutations commonly occur in cancer-associated genes, show evidence of selection and clonal expansion, and may play important roles in tissue regeneration and tumor suppression (Colom et al., 2020; Martincorena et al., 2018, 2017, 2015; Zhu et al., 2019). Collectively, these studies indicate a growing realization that somatic mutagenesis and clonal dynamics are likely an important determinant of human health during aging. While the accumulation of somatic mutations in the mitochondrial genome (mtDNA) with age has long been documented, the specific nature of their occurrence, and the consequences for aging, have remained unclear (Review (Sanchez-Contreras and Kennedy, 2022)).

In vertebrates, mtDNA is a maternally inherited ~ 16-17kb circular DNA molecule encoding 37 genes: 13 essential polypeptides of the electron transport chain (ETC), two ribosomal RNA genes, and 22 tRNAs. Mitochondria are involved in a broad range of crucial processes, including ATP generation via oxidative phosphorylation (OXPHOS), calcium homeostasis, iron-sulfur cluster biogenesis, regulation of apoptosis, and the biosynthesis of a wide variety of small molecules (Kowaltowski, 2000). These processes rely on mitochondria such that disruption of the genetic information encoded in mtDNA by mutation leads to dysfunction of these important processes and subsequently induces disease (Wallace, 1999). Unlike nDNA, mtDNA replication is largely independent of the cell cycle. The higher level of mitochondrial genome replication, the absence of several cellular DNA repair pathways, and the lack of protection from histones results in mtDNA mutation rates ~ 100-1000X higher than that of nDNA (Marcelino and Thilly, 1999). Moreover, due to the coding density of mtDNA being higher than nDNA (~ 91% vs ~ 1%), the probability that a mutation disrupts mitochondrial function is greater.

Observational studies have shown that the genetic instability of mtDNA in somatic cells is a fundamental phenotype of aging and may be involved in the pathogenesis of several diseases (Reviewed in (Larsson, 2010)). Collectively, studies examining endogenous mtDNA mutations have shown low levels of G→T/C→A mutations and a preponderance of G→A/C→T and T→C/A→G transitions. This has been interpreted as being contrary to free radical theories of aging by suggesting that reactive oxygen species (ROS) are not the primary driver of mutagenesis in mtDNA (Arbeithuber et al., 2020; Ju et al., 2014; Kennedy et al., 2013; Williams et al., 2013; Zheng et al., 2006). Other notable patterns include an over-abundance of mutations in the mitochondrial Control Region (mCR), an unusual strand bias, a mutational gradient in transition mutations, and a unique trinucleotide mutational signature (Ju et al., 2014; Kennedy et al., 2013; Sanchez-Contreras et al., 2021; Wei et al., 2019). However, while the presence of somatic mtDNA mutations is well documented, a clear causative role in aging remains controversial (Reviewed in (Sanchez-Contreras and Kennedy, 2022)).

One reason for this controversy stems from a poor understanding of when, where, and how somatic mtDNA mutations arise during the normal aging process. Most conclusions regarding the accumulation of mtDNA mutations during aging are based on a limited number of experimental models and tissue types, with data largely focused on brain and muscle due to their perceived sensitivity to mitochondrial dysfunction. Only a small number of pan-tissue surveys have been performed (Li et al., 2021; Ma et al., 2018; Samuels et al., 2013). Importantly, most of these prior studies made use of either “clone and sequence” or conventional next-generation sequencing (NGS) to detect mutations. These approaches are technically limited in their ability to detect heteroplasmy below a variant allele fraction (VAF) of 1-2% (Reviewed in (Salk et al., 2018)). The advent of ultra-high accuracy sequencing methods has shown that most heteroplasmies are present far below this analytical threshold (Arbeithuber et al., 2020; Kennedy et al., 2013). As such, determining the burden of somatic mtDNA mutations in the context of normal aging lags well behind the efforts focused on the nDNA. This is especially pertinent given the heterogeneous nature of tissue decline during aging.

Like the nDNA, somatic mutations in mtDNA have been proposed to be under selection (Suen et al., 2010). Cells have evolved several mitochondrial quality control pathways such as removal of damaged mitochondria by mitophagy, and fusion/fission to maintain a healthy mitochondrial pool (Youle and Narendra, 2011). The formation and expression of deleterious mtDNA mutations is hypothesized to lead to a loss of mitochondrial membrane potential, mitochondrial dysfunction, and induction of mitophagy. This is a potential mechanism by which cells prevent mtDNA mutations from reaching a phenotypic threshold capable of altering cell homeostasis (Rossignol et al., 2003, 1999). Evidence for involvement of quality control machinery in removing somatic mtDNA mutations has been contradictory, with some indicating a clear role for mitophagy and/or fission/fusion, while other evidence indicates no effect (Chen et al., 2010, 2015; Pickrell et al., 2015; Suen et al., 2010). Thus, the role, if any, of the mitochondrial quality control pathways in targeting mtDNA mutations for removal remains unclear.

We and others have previously identified a mitochondrially targeted synthetic peptide, Elamipretide (ELAM; previously referred to as SS-31 and Bendavia), and the NADH precursor nicotinamide mononucleotide (NMN) as interventions that restore mitochondrial function and tissue homeostasis late in life (Reviewed in (Yoshino et al., 2018) and (Obi et al., 2022)). The specific mechanism(s) by which these two compounds ameliorate age-related mitochondrial dysfunction differ. ELAM interacts directly with the inner mitochondrial membrane and membrane associated proteins, stabilizing the mitochondrial ultrastructure and influencing cardiolipin-dependent protein interactions to improve ETC function leading to reduced oxidant production, preservation of membrane potential, and enhanced ATP production (Campbell et al., 2019; Mitchell et al., 2020; Zhang et al., 2020). In contrast, NMN is an NAD+ precursor molecule and acts by elevating NAD levels and providing additional substrate for mitochondrial ATP generation(Guan et al., 2017; Martin et al., 2017; Yoshino et al., 2011). Neither intervention is expected to directly alter mtDNA repair mechanisms. Therefore, we sought to test whether these interventions would reduce the prevalence of mtDNA mutations in aged tissues because of their proven efficacy in improving mitochondrial structure and/or function.

We first addressed the relative dearth of high accuracy data regarding age-related accumulation of mtDNA in mice across multiple tissue types. To that end, we used ultra-accurate Duplex Sequencing (Duplex-Seq) to identify organ-specific mtDNA mutation burden in heart, skeletal muscle, eye, kidney, liver, and brain in naturally aged mice (Kennedy et al., 2014; Schmitt et al., 2012). Intra-animal comparison allowed us to determine whether mtDNA mutation rates differ between organs while still accounting for inter-animal variation. Our findings point to the accumulation of somatic mtDNA mutations being a dynamic and highly tissue-specific process that can be modulated by one or more cellular pathways amenable to small molecule intervention.

## METHODS

### Animals and tissue collection

C57BL/6j male mice from the National Institute of Aging Rodent Resource were handled according to the guidelines of the Institutional Animal Care Committee at the University of Washington. Two age cohorts were used at 4.5 months and 26 months of age, respectively. Tissues from the 26 month old cohort, including aged mice treated with ELAM or NMN, were obtained from the same previously reported study, as previously described (Whitson et al., 2020). Briefly, 24-month-old mice were randomly assigned to control, ELAM or NMN treatment groups. ELAM was provided by Stealth BioTherapeutics (Newton, MA) and administered at a 3 mg/kg body weight/day dosage for 8 weeks through subcutaneously implanted osmotic minipumps (ALZET, Cupertino, CA). Control mice were simultaneously housed in cages with ELAM pump mice. NMN was obtained from the Imai laboratory (Washington University in St. Louis, MO) and administered through *ad libitum* drinking water with a concentration based on each cage’s measured water consumption and mean mouse body weight to approximate a 300 mg/kg /day dose. This method of drug delivery necessitated that NMN treated mice were housed independently from control animals, however, treatments were run concurrently. Because we sequenced just a subset of the animals from the Whitson *et al*. study (N=3-5 vs N=11-15 each group), we minimized study variation by excluding tissues from animals with clearly cancerous lesions by gross analysis (primarily seen in liver) and then sequenced a random cohort of age matched groups from the three treatment cohorts (Whitson et al., 2020).

Mouse tissue was collected at 26-months of age immediately following euthanasia. Representative portions of six different organ systems were flash frozen: 1) apex of the heart; 2) 2 mm section from the inferior pole of the left lateral liver lobe; 3) Eyes were enucleated and cleared of muscle and adipose tissue before dissecting the retina from the retinal pigmented epithelium (RPE)-choroid complex (also referred to as ‘eye cup’ or ‘EC’ in our raw data files) with both regions preserved separately; 4) 3 mm slice of the lower pole of the decapsulated left kidney; 5) proximal 3 mm of left gastrocnemius; 6) brain was dissected in ice cold 1x PBS to obtain a 3 mm-thick coronal section from the most anterior/septal pole of the left hippocampus and a 3 mm-thick sagittal section from the medial side of the left cerebellar hemisphere. For every sample, dissecting tools were wiped in 70% ethanol, a new razor blades and cutting boards were used, and samples were rinsed in fresh 1x PBS to minimize the contribution of blood and avoid DNA cross-contamination. For perfused experiments, a separate cohort of NIA male mice matching the same age for the aged cohort above (26 mo, N=3) was perfused transcardially with 1x PBS containing calcium and magnesium before tissue isolation.

### DNA processing and Duplex Sequencing

DNA was extracted using the Qiagen DNeasy Blood & Tissue kit (Qiagen, Valencia, CA) and stored at −80°C. Duplex-Seq was performed as previously described (Kennedy et al., 2014), but with several modifications also previously described (Hoekstra et al., 2016; Sanchez-Contreras et al., 2021). Duplex-Seq adapters with defined UMIs were used and were constructed by separately annealing complementary oligonucleotides (IDT, Coralville IA), each containing one of 96 unique molecular identifiers (UMIs) of defined sequence (Table S1). The resulting adapters were diluted to 25μM for ligation to sheared DNA. Targeted capture used the IDT xGen Lockdown protocol and probes specific for mouse mtDNA (Integrated DNA Technologies, Coralville, IA) following the manufacturer’s instructions. The resulting libraries were indexed and sequenced using ~ 150–cycle paired-end reads (300-cycles total) on an Illumina NovaSeq6000 with ~ 20×10^6^ reads per sample. Per sample sequencing metrics are available in Table S2.

### mtDNA content

mtDNA copy number was determined by droplet-digital PCR (ddPCR) by the Genomic Sciences Core (GSC) of the Oklahoma Nathan Shock Center. Briefly, 200ng of total genomic DNA from the same isolated DNA sample used for Duplex-Seq was mixed with ddPCR assay components including fluorogenic ‘TaqMan’ primer probe sets and the reactions were distributed across a chip with ~ 20,000 856pL droplets to dilute the DNA template to either zero or one copy per well, as described previously (Masser et al., 2016). Reactions were then cycled to end-point and fluorescence was read in each droplet. Based on the count of fluorescent positive and negative wells and using a Poisson distribution, the number of target copies was calculated per microliter. Nuclear genome counting was performed in parallel and used as a surrogate for cell number which allows for normalization to cell number and results in an absolute quantitation of mitochondrial genomes. The data are available in Table S3.

### Data analysis and statistics

The raw sequencing data was processed using version 1.1.4 of our in-house bioinformatics pipeline (https://github.com/Kennedy-Lab-UW/Duplex-Seq-Pipeline) with the default consensus making parameters. A detailed description of the Duplex Sequencing pipeline is described in Sanchez-Contreras *et al*. (Sanchez-Contreras et al., 2021). Seven small polynucleotide repeats were masked to reduce errors associated with alignment artifacts (Table S4).

To quantify the frequency of *de novo* events, we used a clonality cutoff of 1% or a depth of <100, which excluded any positions with variants occurring at a high heteroplasmy level and scores each type of mutation only once at each genome position. Called variants were annotated using the Ensembl Variant Effect Predictor v99 to obtain protein change. Mutation frequencies were calculated by dividing the number of reads for each allele by the total number of reads at the same mtDNA position. Correlation statistics were applied to determine intra- and inter-animal mutation frequency variation using GraphPad Prism, R, and Python software.

Statistical significances between young and old mutation frequencies for single nucleotide variants (SNV) and insertions/deletions (In/Dels) were determined by Welch’s t-test for each mutation class between young vs. old. For comparisons within each age group and mutation type but between tissues (e.g. young kidney vs. young liver vs. young heart frequency, etc), one-way repeated measures (i.e. each tissue) ANOVA were followed with Tukey’s HSD For comparing morethan one group to a control (e.g. frequency of mutation type for kidney in aged ELAM or aged NMN treated mice vs. untreated aged control) one-way ANOVA was followed by Dunnett’s multiple comparison test. The significance of the ratio of means between young and old mutation spectra was determined by t-test for the ratio of means of two independent samples from two gaussian distributions with the 95% confidence interval estimated by Fieller’s theorem implemented in the *mratios* R-package (Fieller, 1954)(Fieller 1954). P-values or adjusted p-values less than 0.05 were considered significant in all cases.

Somatic heteroplasmic clones were defined as variants called in >2 supporting reads. Expected clone events were calculated as the percentage of clonal mutations (>2 calls per sample) for each mutation type observed by age, either in total across all samples or by individual tissue types (as indicated). Significance between expected and observed clone events was calculated by Poisson distribution.

dN/dS analysis was carried out using the *dNdScv* R package (Martincorena et al., 2017). The package is designed to quantify selection in somatic evolution by implementing maximum-likelihood methods that accounts for trinucleotide context-dependent substitution models, which is highly biased in mtDNA. Each sample was processed independently using the *dNdScv* implementation with options *max_muts_per_gene_per_sample* set to *Inf, numcode* set to 2, and the mean depth per gene included as a covariate. ND6 was analyzed separately due to it residing on the opposite strand from the other protein coding genes and having different G/C-skew (Ju et al., 2014). The resulting dN/dS values (w_mis_) for each gene were averaged and significant deviation from 1 determined by a one sample t-test with Bonferroni correction (significance set to p≤0.0167).

## RESULTS

To study the effects of aging on the accumulation of somatic mtDNA mutations across tissues, we used Duplex-Seq to obtain high-accuracy variant information across the entire mitochondrial genome.We examined six different organ systems (heart, kidney, liver, skeletal muscle, brain, and eye) at two different ages (young=4.5 months; N=5 and old=26 months; N=6). These two age groups were chosen for their representation of the two extremes of the adult mouse lifespan while mitigating potential confounders related to development, sexual maturation, and survival selection at more advanced ages. These tissues vary on their dependence of mitochondria function and OXPHOS (Fernández-Vizarra et al., 2011). To minimize variation of cell type substructure within tissues between animals, care was taken to isolate similar regions of each organ, as described above. In total, we sequenced over 27.9 billion high-accuracy bases, corresponding to a grand mean post-consensus depth of 10,125X for all samples with reasonably uniform coverage among experimental groups and mice, with the exception of the Ori_L_ (5160-5191) and several masked regions with high G/C content and/or repetitive sequences (Figure S1 and S2; Table S4). We observed a combined total of 77,017 single-nucleotide variants (SNVs) and 12,031 insertion/deletions (In/Dels) across all tissue, age, and intervention groups. Collectively, these data represent the largest collection of somatic mtDNA mutations obtained in a single study to date. A summary of the data for each sample is reported in Table S2.

### Frequency of somatic mtDNA mutations increase with age and is tissue-specific

To better understand the effects of aging on somatic mtDNA mutations across tissues, we determined the frequency of both SNVs and small InDels (≲15bp) in aged mice. To minimize the contribution of mtDNA mutations that could be either maternally inherited or clonal expansions established in development, we limited our analysis to mutations occurring at or below a variant allele fraction (VAF) of 1%. In young mice, an initial comparison of the frequency of mtDNA SNVs revealed a mutation frequency on the order of ~ 1×10^−6^, with low variability between tissues (Figure 1A). Kidney and liver were notable exceptions, exhibiting significantly higher SNV frequencies compared the other tissues in the young cohort (Figure 1A, B). With age, we observed significant increases in SNV frequency in all tissues we surveyed (Figure 1A). Moreover, mutation frequencies varied considerably between tissues in the aged cohort, with kidney having the highest SNV frequency (6.60±0.56×10^−6^) and heart having the lowest (1.74±0.16×10^−6^) (Figure 1A, C). The observed changes in frequency with age or tissue type did not correlate with differences in mtDNA copy number, as the mtDNA:nDNA ratio did not change with age (Table S3; Figure S3A, B). In/Dels were approximately 10-fold less prevalent than SNVs in young mice, with a mean frequency of ~ 1.5×10^−7^, and virtually no differences between tissues (Figure 1D, E). Like SNVs, In/Dels increased with age in nearly all tissues we surveyed and did not correlate with copy number, but unlike SNVs, they did not significantly differ between tissue types, likely due to the high variability between samples (Figure 1D, F; Figure S3C).

**Figure 1.**
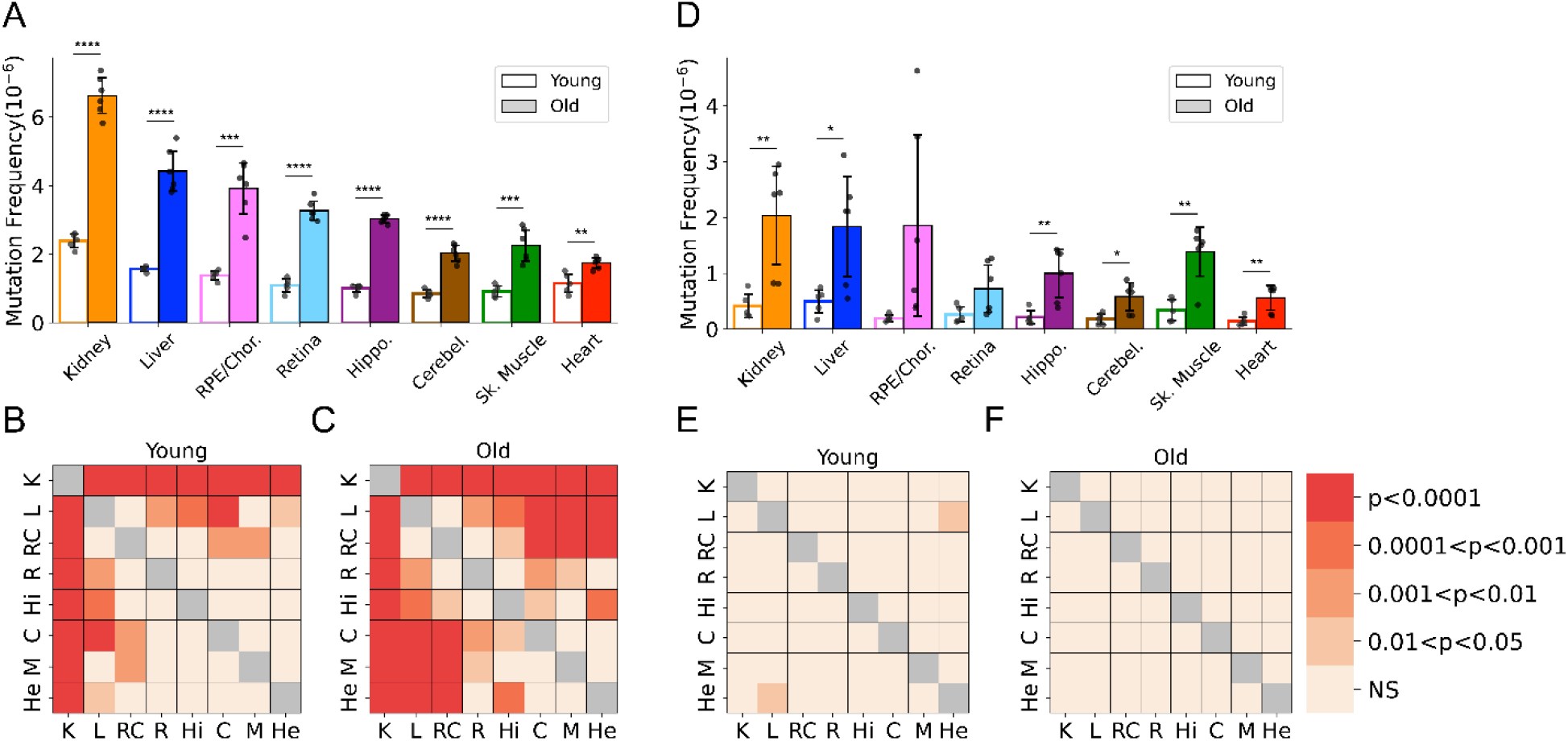
Frequency of somatic mtDNA mutations increase with age and is tissue-specific. **(A)** The frequency by which single nucleotide variants (SNV) were detected in all sequenced bases in either young (~ 5-months-old) or old (26-months-old) tissues arranged from highest to lowest SNV frequency in aged mice. **(B)** The frequency by which DNA insertions or deletions (In/Del) of any size are detected within all sequenced bases either young (~ 5-months-old) or old (26-months-old) tissues. For (A) and (B), significance between young and old within a tissue was determined by Welch’s t-test. *0.01<p<0.05, **0.001<p<0.01, ***0.0001<p<0.001, ****p<0.0001; error bars = standard deviation of individual data points shown. **(C-D)** Heatmaps of one-way ANOVA analysis for significant differences of SNV frequencies between tissues, within either young **(C)** or old **(D)** age groups. **(E-F)** Heatmaps of one-way ANOVA analysis for significant differences of In/Del frequencies between tissues, within either young **(E)** or old **(F)** age groups.

Due to mutation burdens being tissue specific, we considered whether these differences could be driven by variation in the contribution of mitochondrial mutations in leukocytes of circulating blood. To determine this, we analyzed Duplex-Seq in a small subset of tissues from aged mice perfused with PBS to remove the blood. Duplex-Seq of mtDNA from blood collected prior to the perfusion showed that in aged mice, the average frequency of SNV in blood was 3.05±0.15×10^−6^, comparable to the frequency detected in aged hippocampus. Comparisons of perfused (no/low blood) to non-perfused tissues from liver, kidney, skeletal muscle, hippocampus and cerebellum (the retina, RPE/choroid, and heart were not sequenced), showed no significant difference in the frequency of SNV mutations (Figure S4). Thus, our mutation profiles are likely driven primarily by organ-specific cell types.

Although little is known about the kinetics of somatic mtDNA mutation accumulation during aging, they have been reported to increase exponentially during aging in mice (Vermulst et al., 2007). Both this study and the prior study by Arbeithuber *et al*. report only two time points each (4.5-months vs. 26-months and 20-days vs. 10-months, respectively), making it impossible to confirm exponential increase in either study (Arbeithuber et al., 2020). However, the combination of our data with Duplex-Seq data with the previously published data by Arebiethuber *et al*. indicates a linear increase in overall mutation frequencies across the lifespan in the three tissue types common to both studies (brain, muscle, and liver). This indicates a likely constant ‘clock-like’ accumulation analogous to what is seen in the nuclear genome (Abascal et al., 2021; Alexandrov et al., 2015; Arbeithuber et al., 2020) (Figure S5). Together, these data demonstrate mtDNA mutations accumulate at tissue-specific rates during aging and indicate use of a single tissue source to draw broad organism level conclusions regarding the interaction between mtDNA mutations and aging is not scientifically supported.

### Mutation spectra of somatic mtDNA mutations demonstrate tissue-specific distribution of mutation types

Previous work by us and others indicates that somatic mtDNA mutations are strongly biased towards transitions (*i*.*e*. G→A/C→T and T→C/A→G), with low levels of transversions (Ameur et al., 2011; Arbeithuber et al., 2020; Ju et al., 2014; Kennedy et al., 2013; Pickrell et al., 2015; Williams et al., 2013). Moreover, due to their low prevalence transversions associated with oxidative lesions (*i*.*e*. G→T/C→A and G→C/C→G) have been largely discounted as contributing to age-associated mtDNA mutagenesis (Arbeithuber et al., 2020; Hoekstra et al., 2016; Itsara et al., 2014; Kauppila et al., 2018; Kennedy et al.,

2013; Zheng et al., 2006). However, these findings are based on a limited number of tissue types, specifically muscle and brain. Given the wide range of SNV frequencies and known metabolic activities of the tissues we assayed, we examined the mutational spectra for each tissue. Our data show that the overall bias towards G→A/C→T transitions remains broadly true for most tissues, but extent of this bias varies considerably, with kidney and heart being the notable extremes (Figure 2A). In agreement with prior studies, a single mutation class, G→A/C→T, is the most abundant mutation type and accounts for more than 50% all mutations in most young tissues (Figure S6). In contrast, ROS-linked G→T/C→A and G →C/C→G mutations exhibited substantial variation in the level mutations between tissues. In the central nervous system (CNS) tissues (hippocampus, cerebellum, retina), G→T/C→A and G→C/C→G, combined, accounted for an average of 23% of the total mutation burden (Retina = 18%, Hippocampus 18%, Cerebellum 33%) (Figure S6). These data are consistent with prior Duplex-Seq based studies that focused on neural tissues (Arbeithuber et al., 2020; Hoekstra et al., 2016; Kennedy et al., 2013). In contrast, skeletal muscle and heart in young animals have a relatively high frequency of ROS-linked mutations, with 43% and 66% of all mutations, respectively, resulting from these two types of mutations. This suggests that ROS is a greater source of mtDNA mutagenesis earlier in life and is tissue dependent (Figure S6).

**Figure 2.**
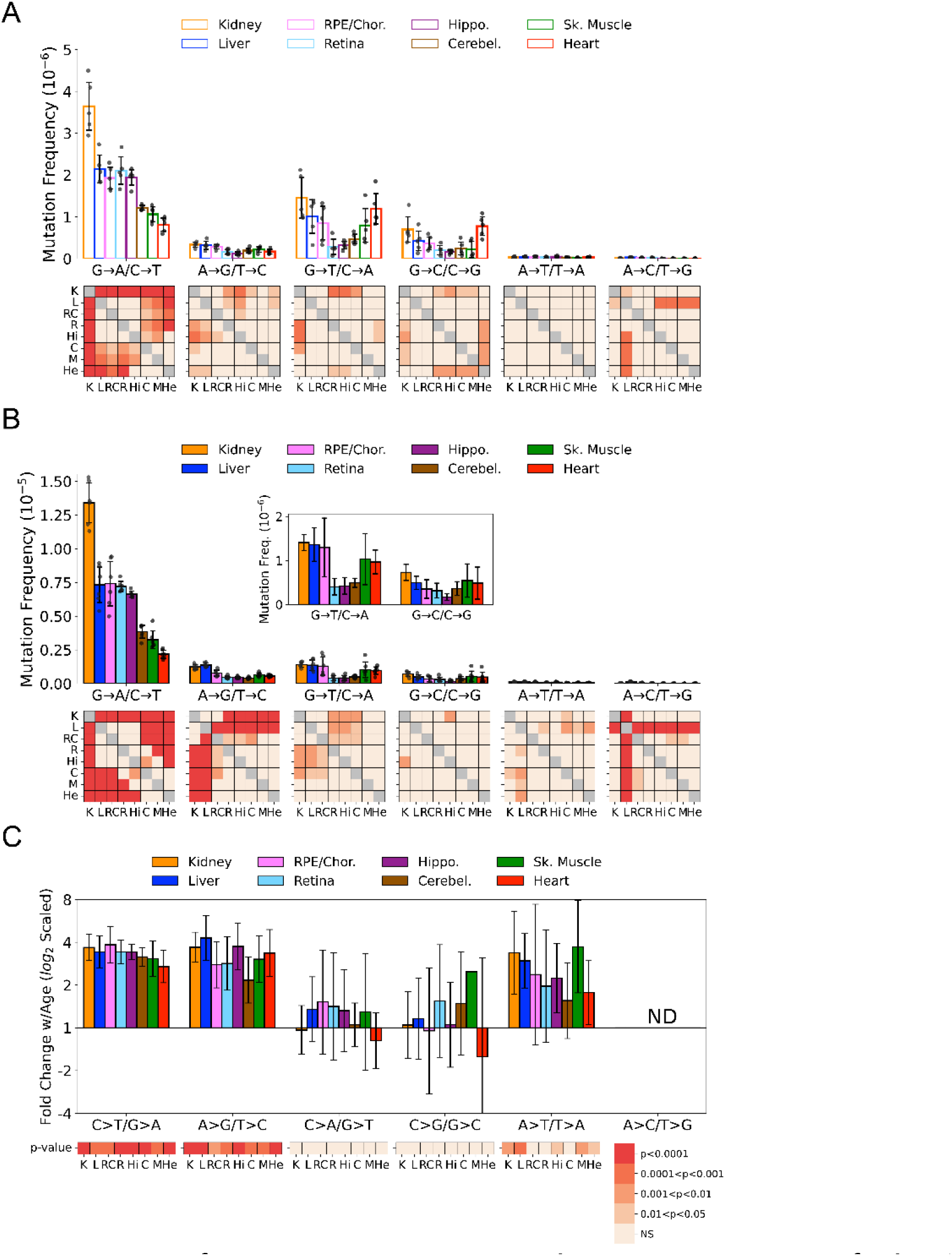
Mutation spectra of somatic mtDNA mutations demonstrate tissue-specific distribution of mutation types. **(A)** SNV frequency by mutation type for young (~ 5-month-old) tissues shows that ROS-linked G→A/C→T mutations largely dictate overall SNV mutation burden and predominate in all young tissues except heart. Tissues of the central nervous system: eye retina, brain hippocampus and brain cerebellum have the lowest frequencies of G→T/C→A and G→C/C→G transversions whereas they are highest in kidney and heart. Heatmaps show p-value from one-way ANOVA analysis for significant differences of SNV frequencies between young tissues within each mutation class. **(B)** SNV frequency by mutation type for old (26-month-old) tissues shows age-specific changes to mutation spectra. Heatmaps show one-way ANOVA for significant differences of SNV frequencies between old tissues within each mutation class. **(C)** Fold change of frequency from young to old age calculated for each tissue and spectra and shown as Log_2_. Heatmap shows whether fold-change values of old relative to young mice are significantly different from fold-change 0 (no change). K = Kidney; L = Liver; RC = RPE/choroid; R = Retina; Hi = Hippocampus; C= Cerebellum; M = Skeletal Muscle; He = Heart.

In comparison to the young tissues, mutation loads became more weighted towards transitions across the aged tissues we surveyed (Figure 2B). Significant differences between tissues within mutation classes also became more pronounced (Figure 2B, *heatmap*). The fold-increase in most mutation types were remarkably uniform despite significant differences in SNV frequency between them (Figure 2C). Aging led to an average 3.2-fold increase of G→A/C→T and T→C/A→G transitions. Similarly, a significant 2.4-fold increase of T→A /A→T transversions was also observed (Figure 2C). A→C /T→G mutations were not evaluated due to their extreme paucity. In contrast to the other mutation types, G→T/C→A and G→ C/C→G mutations did not significantly increase with age (Figure 2C). Thus, we confirmed that ROS-linked mutations do not accumulate significantly with age.

### Clonal expansion of somatic mtDNA mutations is tissue specific

Mutagenesis has been described as an irreversible process that results in increasing levels of mutations in a population over time, termed ‘Muller’s ratchet’ (Felsenstein, 1974; Muller, 1964). Consequently, absent any compensatory mechanisms, mutations should increase during life. In the case of mtDNA, this should appear as an increase in the burden of apparent heteroplasmies (or clones) within a tissue over time. Importantly, because mtDNA replicates independently of nDNA, apparent heteroplasmies can increase during aging even in the absence of substantial cell proliferation. Moreover, mitochondria are subject to surveillance by mitophagy, which may affect the age dependent mutational dynamics in tissue specific ways (Pickles et al., 2018). Expansion of mtDNA mutations sufficient to warrant the term ‘clonal’ has been documented in human tissues, but the prevalence of this phenomenon remains poorly documented in mice (Greaves et al., 2014, 2010, 2006; Nekhaeva et al., 2002). The set of tissues we examined comprise a range of varying proliferative and replicative potentials, with heart, brain and retina being limited, while many kidney and liver cell types proliferate signficantly throughout life. Therefore, we sought to determine the tissue-specific burden and dynamics of age-related clonal expansion of mtDNA mutations.

We defined a “heteroplasmic clone” as a variant supported by three or more error-corrected reads and then calculated both the frequency and percentage of total mutations corresponding to these clones. We observed considerable tissue-specific variation in the effects of age on the presence heteroplasmic clones, with all tissues exhibiting a significant increase in the frequency of total heteroplasmic clones with age (Figure 3A, C). Clones in all tissues were distributed relatively uniformly across the mtDNA but with a striking clustering of variants in the mCR (Figure 3C, *green region*), consistent with prior reports (Arbeithuber et al., 2020; Kennedy et al., 2013; Sanchez-Contreras et al., 2021).

**Figure 3.**
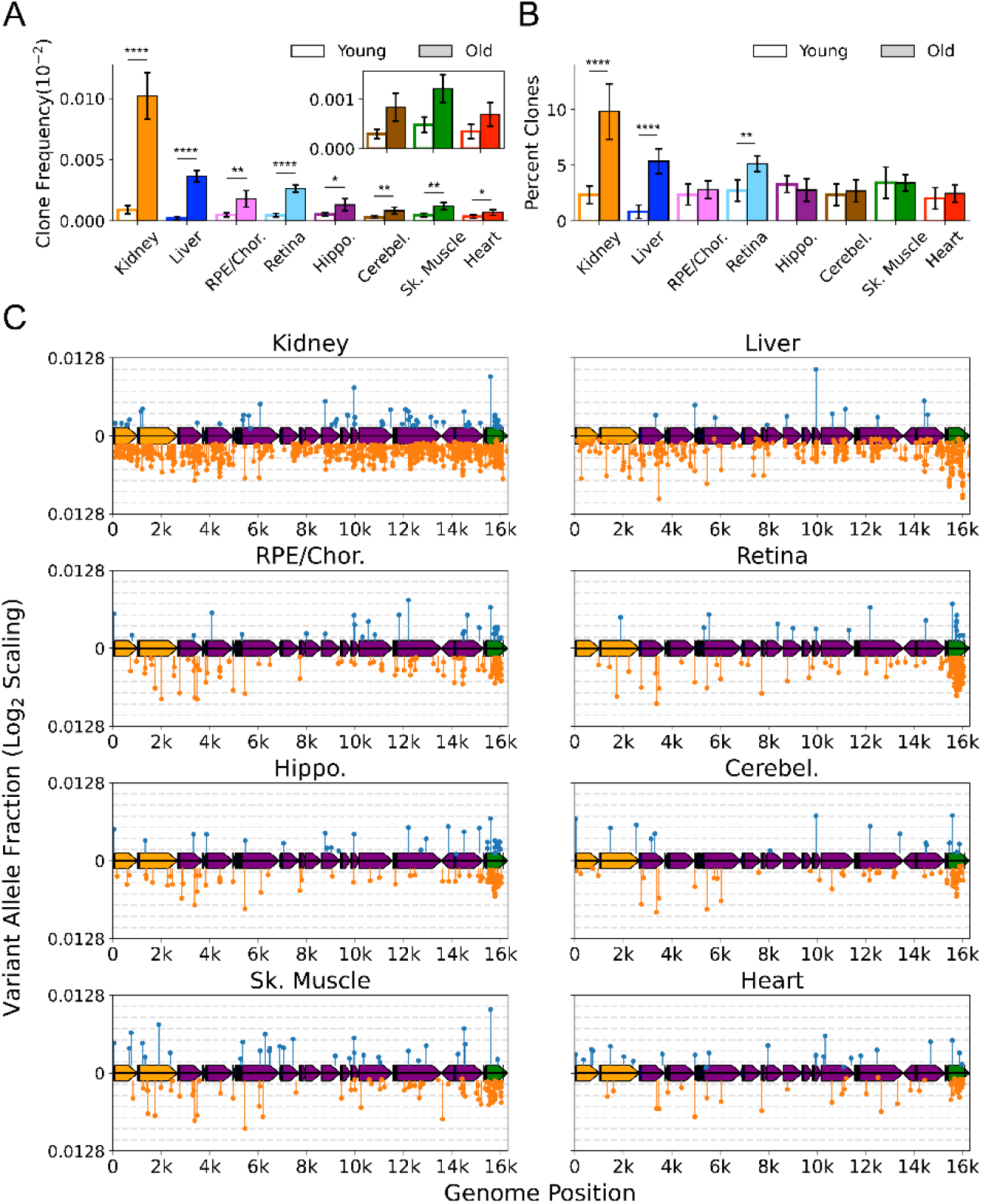
Clonal expansion of somatic mtDNA mutations is tissue specific. **(A)** Frequency of mtDNA clones detected in each tissue shows an increase in detection of clones with age in all tissues. Note that y-axes are set for each tissue (n=5 for young; n =6 for old, error = +/-standard deviation). **(B)** Percentage of total mutations found in heteroplasmic clones for each tissues shows that only kidney, liver and retina have significant increases in relative ‘clonality’ with age. For (A) and (B), significance between young and old within a tissue was determined by Welch’s t-test. *0.01<p<0.05, **0.001<p<0.01, ***0.0001<p<0.001, ****p<0.0001; error bars = standard deviation of individual data points shown. **(C)** Lollipop plots show the mtDNA genomic location of clonal hetroplasmic mutations in young (top row, blue markers, n=5) and old (bottom row, orange markers, n=6) for each tissue type. Orange =rRNA; Dark Blue = tRNA; Purple = Protein Coding; Green = Ori_L_ or mCR.

The mutation composition of the clones varied between tissues. In the RPE/choroid, brain, skeletal muscle and heart, the relative percentage of SNVs found as heteroplasmic clones did not change with age (~ 2-3% of total SNV). In contrast, kidney, liver, and retina exhibited a disproportionate increase in the number of clonally expanded variants with age. In old kidney, the percentage of SNV detected as heteroplasmic clones increased to 10.4%, while clones in liver increased from ~ 1% of SNVs in young to 5.6% in old (Figure 3B). In retina, the percentage of SNVs detected as clones increased from 3.6% in young to 5.6% in old mice. In kidney and liver, the expansion of mtDNA mutations was pervasive across the genome and suggestive of a relationship to the high proliferative and regenerative capacity of these tissues. Retina, however, is a post-mitotic tissue and displayed a very different pattern, with the age-associated increase in clonality being attributed almost entirely to variants clustered in the mCR (Figure 3C). Importantly, several of the tissues we examined are highly vascular and the hematological compartment has been documented to exhibit significant heteroplasmy and shifts in clonal expansions with age (Lareau et al., 2019). With the exception of kidney, which showed a modest effect, we observed no significant changes between the perfused and non-perfused samples, indicating that blood is not a significant source of age-dependent changes across tissues (Figure S7). Collectively, these data suggest that the importance of mtDNA heteroplasmic clones in aging phenotypes is tissue dependent. Very few studies have examined somatic mtDNA heteroplasmic clones in any tissue, and this remains an area for future work.

### Spectral analysis of clonal somatic mtDNA mutations suggests removal of ROS-linked mutations

Having established the tissue-specific profile of heteroplasmic clones, we reasoned that we could distinguish between mutations arising from a transient process earlier in life and an active clearance of mtDNA and/or whole mitochondria containing mutation types by examining which mutation types became more heteroplasmic with age. Specifically, an ongoing or early transient mutational process would be expected to result in expansion of a subset of variants across all mutation types. In contrast, evidence of active clearance would appear as either a lack of heteroplasmic expansion or a bias in the specific variants that underwent expansion. Because five of the eight tissues showed no significant change in the proportion of clonal SNV with aging, we expected that clones would be distributed across the spectra in a pattern like that of non-clonal mutations. Instead, we observed that the spectrum of heteroplasmic clones demonstrates a nearly complete suppression of clones derived from G→T/C→A and G→C/C→G mutations in both young and old mice (Figure 4A, B). Remarkably, suppression of ROS-linked clones was true even in the heart, which carried the highest combined G→T/C→A and G→C/C→G SNV mutation burden of any tissue, with 65% of total SNV in young and 34% of SNV in old mice (Figure S6). Thus, we asked whether this lack of G→T/C→A and G→C/C→G clones was a significant finding or merely a consequence of low sampling due to the relative paucity of clones and the lower frequency of ROS-associated mutations. Under the assumption that clones arise randomly, we tested whether our data set of detected SNV clones differed from the expected number of clones based on the spectral distribution of non-clonal SNV mutations. To ensure that we had sufficient power, we combined SNV mutations and clones detected in all eight tissues of the six old mice for a total of 24,244 *de novo* SNVs and 1,461 heteroplasmic clones. To account for differences in depth between samples, the spectral distribution of total SNV mutations was calculated for each tissue of each mouse. The expected contribution of clones in each tissue was then weighted based on the average percent of ‘clonality’ measured in the aged data set (Figure 4B). For example, more clones were expected to form in kidney and liver (10.4% and 6% clonality, respectively) than would be expected in brain or muscle tissues (~ 3% clonality). Using this method, our model predicted that we should expect to detect 1,341 clones in total for combined aged tissues, which was within 10% of the detected clone total of 1,461.

**Figure 4.**
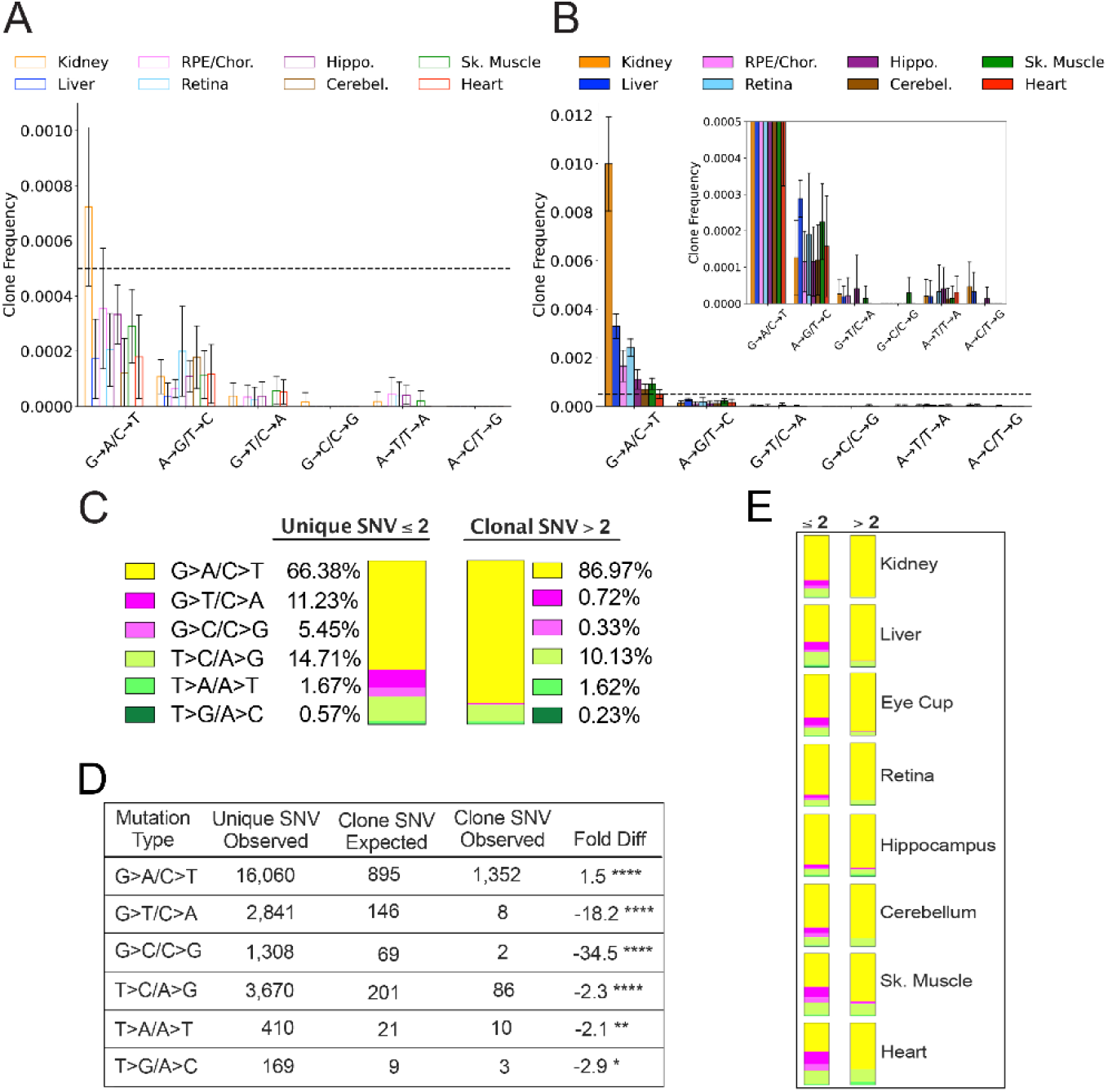
Spectral analysis of clonal somatic mtDNA mutations suggests removal of ROS-linked mutations. **(A)** Frequency of heteroplasmic clones in young mice shown as clone frequency for each mutation class and tissue. **(B)** Frequency of heteroplasmic clones in aged mice shown as mutations per genome (MPG) for each mutation class and tissue. Inset shows graph with adjusted axis to match young mice in (A) to better visualize lack of mutations in G→T/C→A and G→C/C→G mutation classes despite expansion of clonality with age. In both (A) and (B) the dotted line indicates frequency value of 0.005. **(C)** The combined distribution of mutation spectra for single nucleotide variants (SNV) for either unique mutations (detected 2 or less times per single sample) or clonal mutations (detected more than 2 times per single sample) for all aged tissues (48 samples in total). **(D)** Table showing that, based on the mutation spectra of unique mutations, the observed number of SNV clones differs the number SNV clones expected if heteroplasmic clones arise randomly as a consequence of mutation burden. G→A/C→T and T→C/A→ G clones are over-represented, while G→T/C→A and G→C/C→G clones are strongly under-represented based on a Poisson distribution. ‘Fold Diff’ represents fold change of observed clone values relative to expected. **(E)** Mutation spectra distributions for each aged tissue type show that clonality within individual tissue types mirrors the pattern of the combined aged samples with over/under representation of specific mutation types within observed clones. ≤2 = unique mutations, <2 = clonal mutations, mutation types are color coded as in (C).

We modeled the expected spectrum of these clones among the six mutation types using a Poisson process to model random sampling error due to the low abundance of clones. We compared the expected number to the observed spectrum and found that the clonal spectra differed significantly from the distribution of non-clonal SNVs (Figure 4C, D). G→T/C→A and C→G/G→C mutations were predicted to form ~ 146 and ~ 69 clones respectively, however, only eight G→T/C→A clones and two C→G/G→C clones were detected in the entire aged mouse data set, corresponding to an 18- and 34-fold underrepresentation, respectively. Conversely, G→A/C→T and T→C/A→G transitions only deviated from the expected values by less than two-fold (Figure 4D). Although we used a combined aged tissue data set to ensure that we were not under sampling, this under/over representation by mutation spectra was detected in every tissue type as shown by their observed spectral distribution (Figure 4E). Taken together, these results suggest that expansions of heteroplasmic clones in mtDNA do not arise as a random consequence of somatic mutation formation. The uneven distribution of clones relative to non-clonal SNVs suggests that the lack of G→T/C→A and G→C/C→G mutation accumulation with age does not reflect differences in the formation of these mutations, but rather is consistent with a steady-state level of ROS-linked mutations that is susceptible to a constant generation and removal.

### Late-life treatment with mitochondrially-targeted interventions eliminate ROS-linked mutations

Like in the germline, mtDNA mutations have been hypothesized to be selectively removed in the somatic tissue through a mechanism involving a still undetermined interaction between the unfolded protein response, mitophagy, and mitochondrial fission/fusion (Chen et al., 2010; Gitschlag et al., 2016; Lin et al., 2016; Suen et al., 2010). Therefore, we hypothesized that compounds known to improve function and/or ultrastructure in aged mitochondria would impact the burden of aging mtDNA mutations by shifting the steady state of ROS damage towards removal of damaged/dysfunctional mitochondria and their accompanying mtDNA. To this end we sequenced tissues from mice treated systemically for eight weeks with either ELAM or NMN in old mice that had accumulated somatic mtDNA mutations throughout their lives. Functionally, both ELAM and NMN, improve mitochondrial energetics and the mitochondrial network in aged mice across multiple aged tissues within an eight-week time frame (Chiao et al., 2020; Sweetwyne et al., 2017; Whitson et al., 2020), albeit through different mechanisms. This allowed us to examine mice within such a narrow window of time that it was more likely to detect changes in mutation turnover/removal, rather than significant prevention of mutation accumulation during aging.

All treated samples were sequenced to a similar mean ‘duplex’ depth and detected comparable numbers of mutations as the controls (Figures S2, S8, and S9; Table S2). We did not observe a significant change in the overall mutation frequencies between aged mice and treated mice, regardless of tissue, indicating that these interventions do not indiscriminately affect mtDNA mutations (Figure S10). In support of this observation, the non-synonymous to synonymous ratio (dN/dS), which is a measure of positive or negative selection, shows no significant deviation from the expected ratio of one for any age, tissue, or intervention (Figure S11). However, consistent with our hypothesis, ELAM significantly reduced G→T/C→A transversions in heart (Control: 9.7±3.0×10^−7^ vs. ELAM: 5.5±1.5×10^−7^, p=0.019; Control: 9.7±3.0×10^−7^ vs. NMN: 4.7±0.9×10^−7^, p=0.017). As well, A→T/T→A mutations were reduced in NMN treated animals relative to control (p=0.0018) (Figure 5A). Kidney, trended lower in ROS-linked mutations in NMN treated animals with reduction in G→C/C→G mutations reaching significance in NMN treated kidneys (Control: 7.3±2.0×10^−7^ vs. NMN: 3.7±1.7×10^−7^, p=0.038, Figure 5B). Liver showed a trending decrease in G→T/C→A mutations with NMN intervention (Control: 1.4±0.4×10^−6^ vs. NMN: 6.3±1.9×10^−7^, p=0.120, Figure 5C). Muscle showed a trending decrease in G→A/C→T mutations with ELAM intervention (Control: 3.3±0.7×10^−6^ vs. NMN: 2.7±0.3×10^−6^, p=0.110 Figure 5C). Because G→T/C→A and G→C/C→G mutations do not increase with age but are reduced with these interventions, these results indicate that these changes are not simply due to the prevention of these mutation types during the treatment window. In contrast, tissues of the central nervous system, including cerebellum, hippocampus, and retina as well as the RPE/choroid did not display the same pattern in G→T/C→A mutations as seen in peripheral tissues, with no reduction in ROS-linked mutations for treated mice (Figure S12). In cerebellum, A→T/T→A mutations trended slightly higher in NMN treated mice compared to controls. These findings are consistent with the lower overall G→T/C→A and G→C/C→G mutation burden in these latter tissues and may indicate that their mtDNA is more protected from ROS damage with age than is the case in peripheral organs.

**Figure 5.**
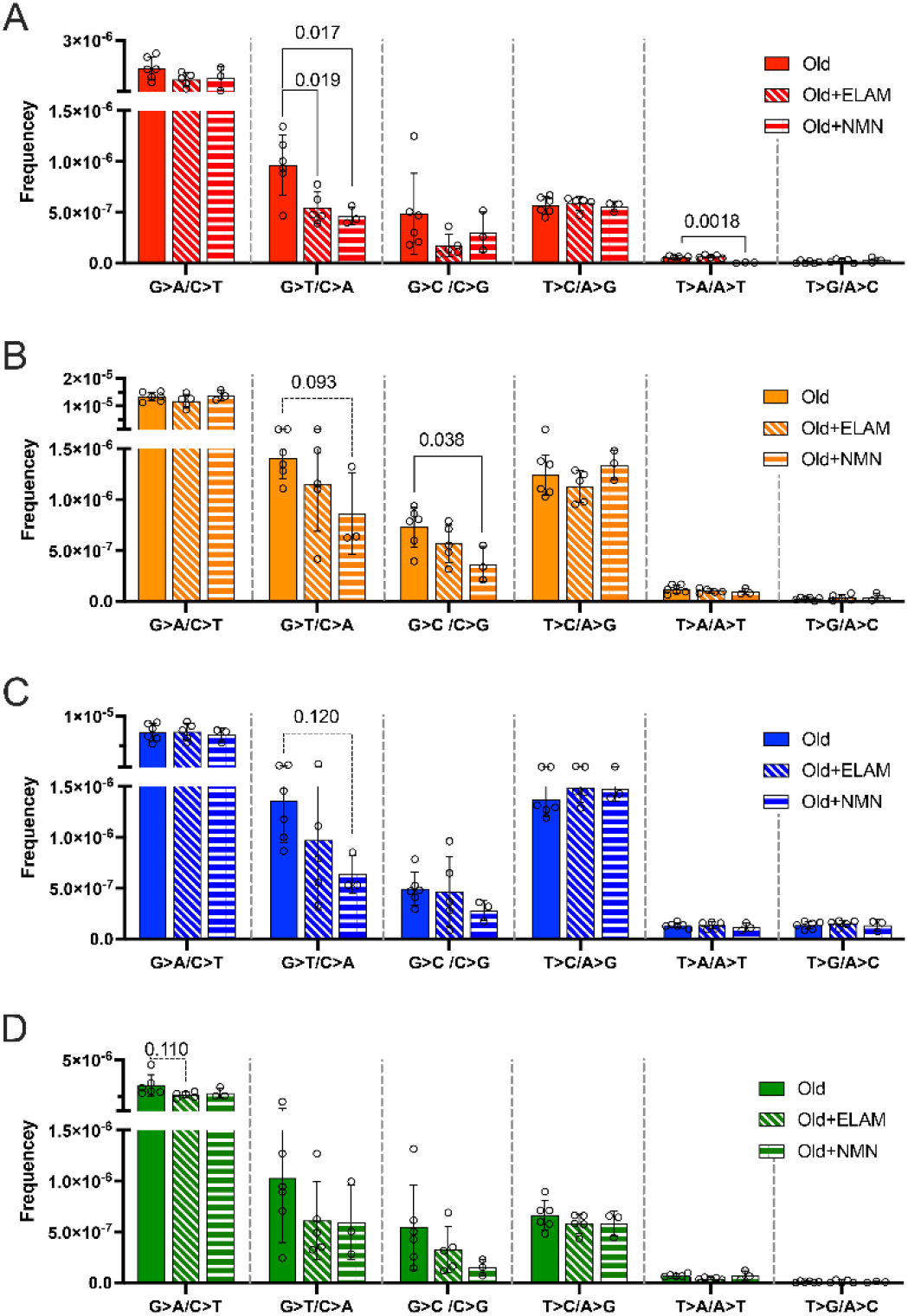
Late-life treatment with mitochondrially-targeted interventions reduces somatic mtDNA mutation burden and is consistent with a mechanism of active ROS-linked mutation removal. Mutation spectra show that aged mice treated for 8 weeks with either elamipretide (ELAM, diagonal striped bars) or nicotinamide mononucleotide (NMN, horizontal striped bars) have decreased frequency of mutations specifically in mutation types linked to oxidative damage, G→T/C→A and G→C/C→G, specifically for **(A)** Heart (red); and **(B)** Kidney (orange) and trending for **(C)** Liver (blue). G→C/C→G mutations are significantly lower in NMN treated kidneys (b). **(D)** Muscle (green) trends lower for ELAM treated mice in G→A/C→T mutations. Error bars = +/-standard deviation. Statistics calculated by one-way ANOVA for each mutation class within tissue, Dunnett’s multiple comparison test compared to untreated aged control group, solid line = significant p<0.05, dotted line = trend p<0.15.

## DISCUSSION

The processes that drive and influence somatic mtDNA mutagenesis in aging and disease has proved to be nuanced and controversial (Kauppila and Stewart, 2015; Sanchez-Contreras and Kennedy, 2022; Szczepanowska and Trifunovic, 2017). Enhancement of accuracy by NGS and newer methods, such as Duplex-Seq, have begun to shed light on these processes. The few studies that have applied these approaches to mtDNA have noted that the distinct lack of G→T/C→A and G→C/C→G transversions, suggesting that oxidative damage is not a major contributor to aging mtDNA mutagenesis (Ameur et al., 2011; Andreazza et al., 2019; Arbeithuber et al., 2020; Hoekstra et al., 2016; Itsara et al., 2014; Kennedy et al., 2013; Samstag et al., 2018; Zheng et al., 2006). While these studies have increased our understanding of mtDNA mutagenesis, these conclusions are largely based on a small number of tissue types. In this report, we took advantage of our continued improvement in the Duplex-Seq protocol to perform a multi-tissue survey of somatic mtDNA mutations in a naturally aging mouse cohort. The very high depths we achieved (depth grand mean: 10,125X) allowed us to detect hundreds to thousands of mutations per sample (variant count mean: 453, Total: 77,017) across all mutation types, representing a ~ 8.3-fold increase in depth and a 38.5-fold increase in mean mutation count per sample compared to the next largest currently published Duplex-Seq dataset for mouse mtDNA (depth grand mean: 1,231X; variant count mean: 12.7, Total: 2,488) (Arbeithuber et al., 2020). This substantially expanded dataset allowed us to examine the types and classes of *de novo* mtDNA mutations with unprecedented sensitivity and as a result, we observed unexpected patterns that would have been missed with a lower coverage of mutations. For example, our data set observed a mean of 50 G→T/C→A mutations per sample with no samples having zero instances. In contrast, Arbeithuber *et al*. reported a mean of three G→T/C→A mutations per sample with ~ 17% of samples failing to detect this mutation type at all (Arbeithuber et al., 2020). Such low numbers can lead to significant biases in determining mutation frequencies and make it difficult to detect meaningful differences between sample types. This issue is exacerbated in the detection of heteroplasmic clones, which account for only a small fraction of detected SNV.

An analogous issue is at play when examining heteroplasmic clone dynamics. In the absence of a significant technical error-rate, such as afforded by Duplex-Seq, the detection threshold of a heteroplasmy is dictated primarily by the depth of sequencing. Therefore, the lower the depth, the larger the VAF must be in order to be detected. Prior studies that made use of conventional NGS to detect low-level mutations in mtDNA required VAF thresholds of 0.5-2% to be above error rate of these platform and reported no changes with age or genotypes being studied (Ameur et al., 2011; Kauppila et al., 2018; Ma et al., 2018). In contrast, recent work by Arbeithuber *et al*., as well as the data reported here, note clear age-related changes occurring well below the detection threshold of earlier studies in mice, suggesting that high detection thresholds imparted by either high error-rates or limited depth do not allow for adequate sensitivity to detect these relatively subtle changes (Arbeithuber et al., 2020). The very high depths achieved in this study also allowed us to observe heteroplasmic clones at a median VAF of 2×10^−4^, which is ~ 8.5-fold lower than the minimum VAF of 1.7×10^−3^ reported in Arbeithuber *et al*. (Arbeithuber et al., 2020). These much lower levels allowed us to detect changes in both absolute and relative burden that would have been difficult to discern with less data or lower accuracy methods, thus highlighting the essential need to use high accuracy methods as the *de facto* approach to testing hypotheses related to somatic mutations.

The findings reported here broadly recapitulate those from smaller studies reporting that somatic mtDNA point mutations occur at a frequency on the order of 10^−6^, increase with age, and are biased towards G→A/C→T transitions (Ameur et al., 2011; Andreazza et al., 2019; Arbeithuber et al., 2020; Hoekstra et al., 2016; Kennedy et al., 2013; Samstag et al., 2018; Williams et al., 2013). However, our expanded dataset found an unexpected level of variation between tissues in both overall mutation frequency and spectrum, indicating tissue-specific effects of aging on mtDNA mutation burden. In young mouse tissues, we observed minimal variation in mtDNA SNV frequency with only kidney, liver, and RPE/choroid showing significantly increased levels compared to other tissues. Only with advancing age were distinct differences between all tissues apparent. Interestingly, although the relative mutation frequencies differed widely between tissues in our data set, we observed a surprisingly consistent two-fold increase from young to old mice, driven largely by the accumulation of G→A/C→T transitions.

Combining our Duplex-Seq data with those from Arbeitheruber *et al*. show a clear clock-like behavior reminiscent of the Horvath epigenetic clock (Horvath, 2013). Such a clock-like phenotype holds promise as a biomarker for biological or even chronological age, depending on its modifiability. Our data showing that small molecule interventions can indeed modify mtDNA mutations suggests that changes in the mutational clock may be possible.

The age-related spectrum of mutations between tissues revealed considerable variation of the canonical ROS-associated G→T/C→A and G→C/C→G transversions (Figure 2). Interestingly, and consistent with prior reports, mouse tissues that are part of the CNS exhibited distinctly reduced levels of ROS-associated transversions compared to the other tissues we surveyed (Figure 2) (Ameur et al., 2011; Kennedy et al., 2013; Williams et al., 2013). Neural tissues are widely considered to be exquisitely sensitive to ETC dysfunction and therefore may have evolved on their ability to defend from it. More robust repair or quality control mechanisms may help eliminate damaged mtDNA before inducing mutagenesis. Consistent with this idea, we observed that both heart and skeletal muscle have a high relative burden of G→T/C→A and G→C/C→G transversions, especially in young tissues, suggesting that their high metabolic activity may confer transversion mutagenesis in these tissue types relative to transitions. These observations suggest that previously broad conclusions of ROS being irrelevant in mtDNA mutagenesis may be biased for having relied on CNS tissues. As shown in previous Duplex-Seq studies, ROS-linked mtDNA mutations are abundant in young, striated muscle tissues but they did not increase in prevalence with age. Contrary to what would be expected, ROS-linked mutations very rarely resulted in expanded heteroplasmic clones regardless of their contribution to overall mutation burden in all tissues we assayed. The disproportionate lack of G→T/C→A and G→C/C→G clones suggests that mutations arising from ROS damage to the mtDNA are cleared more often than mutations formed during replication.

The data reported in this study indicate that mtDNA mutations accumulated very differently across tissues. Mutations occurring from replication are likely to arise with replication in any genome, even in fully functional mitochondria. However, there are some aspects of mtDNA structure and replication patterns that prevent polymerase gamma (PolG)-induced mutations from being distributed entirely randomly across the genome. First, we have previously used the power of this large data set to demonstrate that a gradient exists across the mtDNA genome in the formation of G→A/C→T mutations, driven by the structure and composition of the genome itself (Sanchez-Contreras et al., 2021). Second, the relative prevalence of these mutations, as well as their increased percentage of clonal mtDNA heteroplasmy in aged kidney and liver tissues, likely demonstrates how cellular proliferation also contributes to the accumulation of mtDNA replication-linked transitions. Despite these caveats, PolG errors likely explains why G→A/T→C mutations appear to be tightly linked with age across all tissues. In contrast, ROS-linked mutations are not specifically generated by the process of DNA replication itself. It was unexpected to find that such a significant proportion of mutations in some young tissues were ROS-linked, including 65% of all SNV mutations in the heart and 43% in skeletal muscle. Despite this, there was little increase of these mutations with age such that the proportion of ROS-linked mutation burden relative to all SNV mutations dropped to 30% and 25% heart and skeletal muscle respectively. This finding suggests the possibility that different tissues experience varying levels of ROS injury in early life, but that these differences are attenuated in aged animals. Alternatively, it could mean that there is tissue-specificity in how cells repair and/or destroy oxidatively damaged mitochondria and/or mtDNA resulting in a steady-state of ROS-linked mutations.

We propose that instead of the incidence and impact of ROS damage on mtDNA being minimal, recognition and removal of ROS-linked mutations are maintained at a steady state during aging. We expect that in tissues with exacerbated levels of ROS, the extent of damage could be sufficient to simultaneously create multiple lesions along an individual genome. This could affect the formation of ROS-linked mutations if enough damage is accrued to targeting that mitochondrion for autophagic removal. Alternatively, mitochondrial dysfunction itself can elicit apoptosis, which would also effectively remove damaged mtDNA from the pool. Finally, if replication is initiated, significant DNA damage can stall progression of the replication fork indefinitely, causing collapse of the replication machinery and leading to large mtDNA deletions. This last scenario is a blind spot in Duplex-Seq methodology, such that large deletions are difficult to detect due to the reliance on alignment of barcoded DNA fragments of ~ 200-400 nucleotides. These scenarios all would prevent ROS-induced DNA damage from being converted into a mtDNA mutation.

Our study is the first to demonstrate that ROS-linked mtDNA mutations can be specifically decreased pharmacologically at late age and within a short treatment period in some tissues. These pharmacological interventions are known to reverse age-related decline in mitochondrial function by improving the function of the mitochondrial pool in some aged tissues, albeit through very different mechanisms. Although neither drug has been shown to interact directly with mtDNA, the reduction of ROS-linked mtDNA mutation frequency within such a short treatment window suggests a measurable loss of mitochondrial genomes harboring ROS-linked mutations, rather than a reduced rate of accumulation due to reduced excess ROS. This response demonstrates why ROS-linked mtDNA mutations are rarely found to accumulate with age. Left unexplored by our current study is whether an earlier initiation, longer duration, or more optimal dose of intervention would more robustly alter the mutation spectra. Additionally, we cannot discount that the lack of efficacy in some tissues may be due to tissue-specific differences in biodistribution of the two compounds tested. Finally, we did not have the opportunity or sufficient sample size in this study to explore potential correlation between the heterogeneity in mitochondrial function of tissues versus the degree of change in the oxidative mutations.

Our data indicate that the accumulation of somatic mtDNA mutation is highly segmental in nature and that its impact on aging and/or disease risk may be tissue dependent. Moreover, several of the tissues we examined, such as kidney, are very heterogeneous in their cell-type composition. By examining two compartments of brain (hippocampus and cerebellum), two compartments of the eye (retina and RPE/choroid), and two types of striated muscle tissue (skeletal and heart) we observe that not only are aging-related mtDNA mutation patterns tissue-specific, but they are also region and cell type-specific. The different mutation spectra that we detected must be underlain by cell-specific regulation of mtDNA. Previous overarching assumptions of how mtDNA mutations do, or do not, contribute to aging may have been premature when based on limited information from few tissues and with previous technical hurdles of poor accuracy in detection of low-level mutation burdens. To fully understand how somatic mutation of mtDNA contributes to common diseases of aging, future work must delve into cell-specific mechanisms of mitochondrial mutation accumulation and mtDNA regulation, combined with high-accuracy methods of mtDNA mutation detection.

## Data Availability and Reproducibility

The Duplex-Seq-Pipeline is written in Python and R, but has dependencies written in other languages. The DuplexSeq-Pipeline software has been tested to run on Linux, Windows WSL1, Windows WSL2 and Apple OSX. The software can be obtained at https://github.com/KennedyLab-UW/Duplex-Seq-Pipeline. Raw mouse sequencing data used in this study are available at SRA accension PRJNA727407. The data from Arbeithuber *et al*. are available at SRA accension PRJNA563921. The final post-processed data, including variant call files, depth information, data summaries, and mutation frequencies, as well as the scripts to perform reproducible production of statistics and figure generation (with the exception of Figure 4C-E) are available at https://github.com/Kennedy-Lab-UW/Sanchez_Contreras_etal_2022.

## Acknowledgements

The authors wish to acknowledge the technical support of Carolyn N. Mann for the Sweetwyne Laboratory and Dr. Claudia Moreno for the helpful discussions and long sessions of scientific writing.

## Author contributions

M.S.C. and M.T.S. conceived of and performed mouse tissue experiments and isolations, DNA extraction and Duplex Sequencing. M.S.C., M.T.S. and S.R.K. analyzed the results and wrote the manuscript. J.A.W., M.D.C. and M.T.S. set up the mouse colonies and organized the tissue collections. K.A.T. collected retina tissue, dissected RPE from retina and contributed to the first draft of the manuscript. B.F.K. and S.R.K. designed and developed computational programs for duplex sequencing data. H.J.K. and M.S.C. developed R scripts to query and analyze data. M.J.H., J.F. and M.N. performed experiments. J.H., D.M and P.S.R. provided financial support and assistance in supervising the project with M.T.S and S.R.K. All authors reviewed the final manuscript.

## Figures and Figure Legends

**Figure S1.**
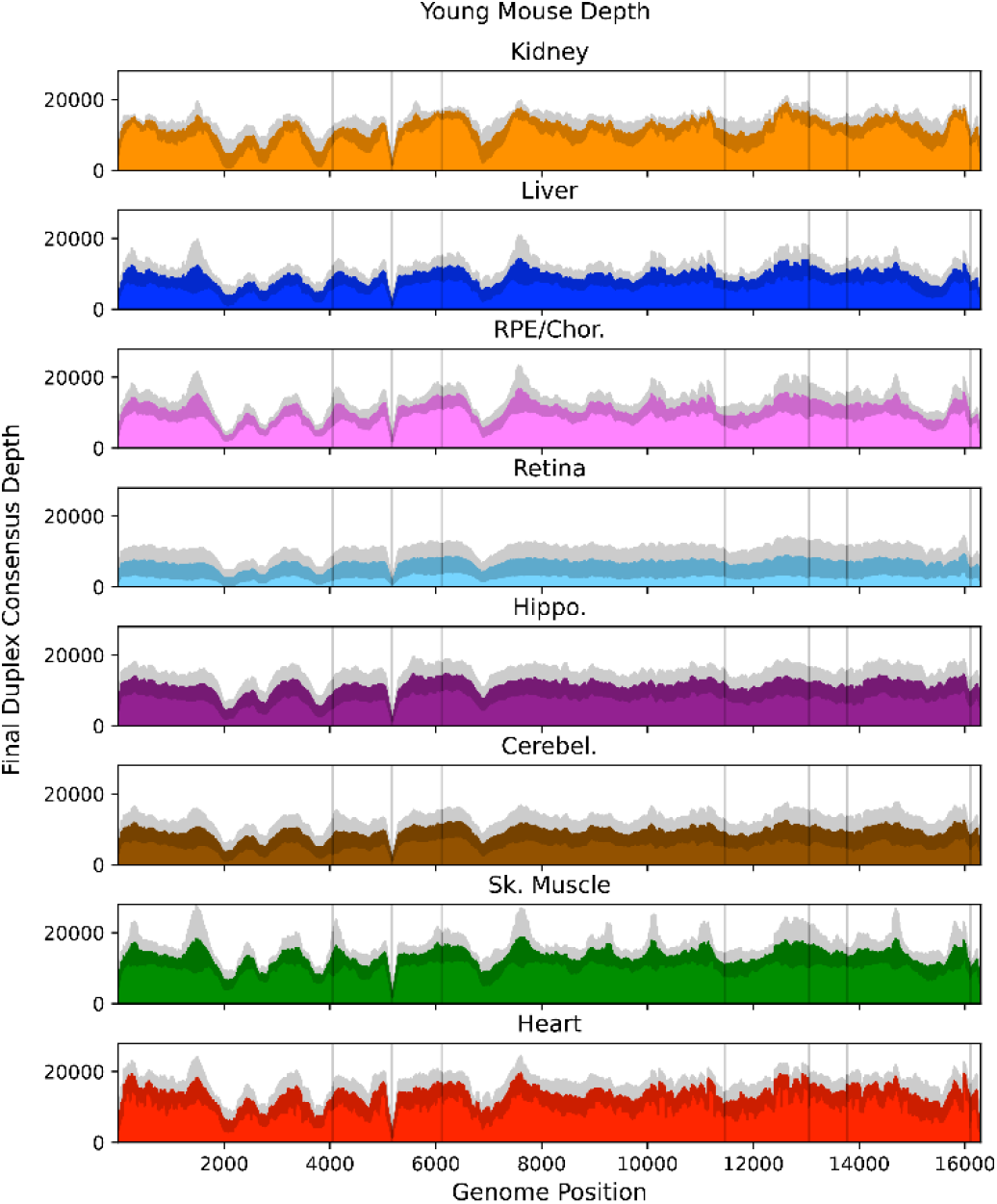
Mean post-consensus ‘Duplex’ depth for young (4.5mo) tissues. Variant occurring within the masked regions (*vertical lines*) or positions with less than a post-consusensus depth of 100X were ignored. *Grey shading* = standard deviation of the mean for N=5 mice.

**Figure S2.**
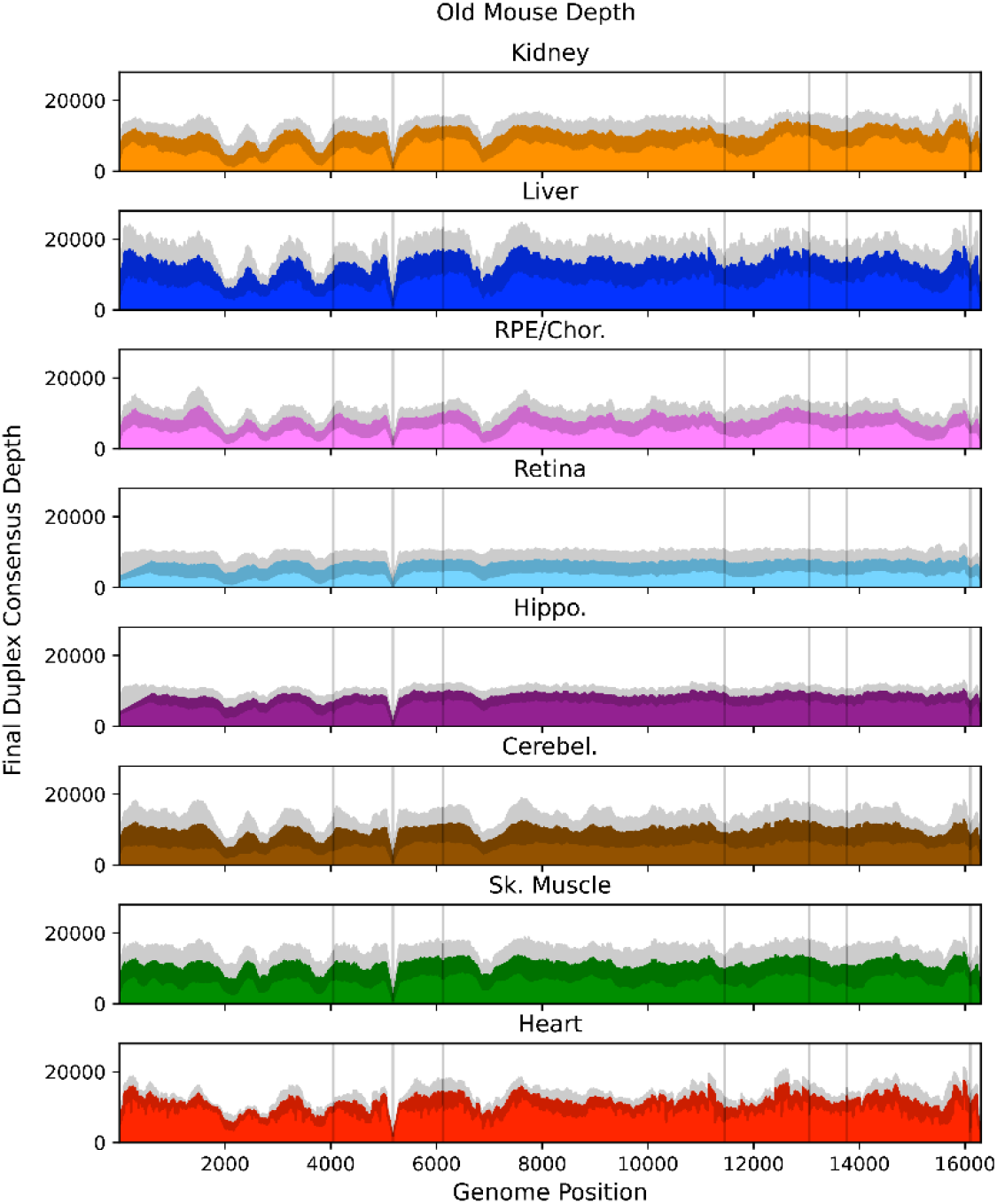
Mean post-consensus ‘Duplex’ depth for old (26 mo) tissues. Variant occurring within the masked regions (*vertical lines*) or positions with less than a post-consusensus depth of 100X were ignored. *Grey shading* = standard deviation of the mean for N=6 mice.

**Figure S3.**
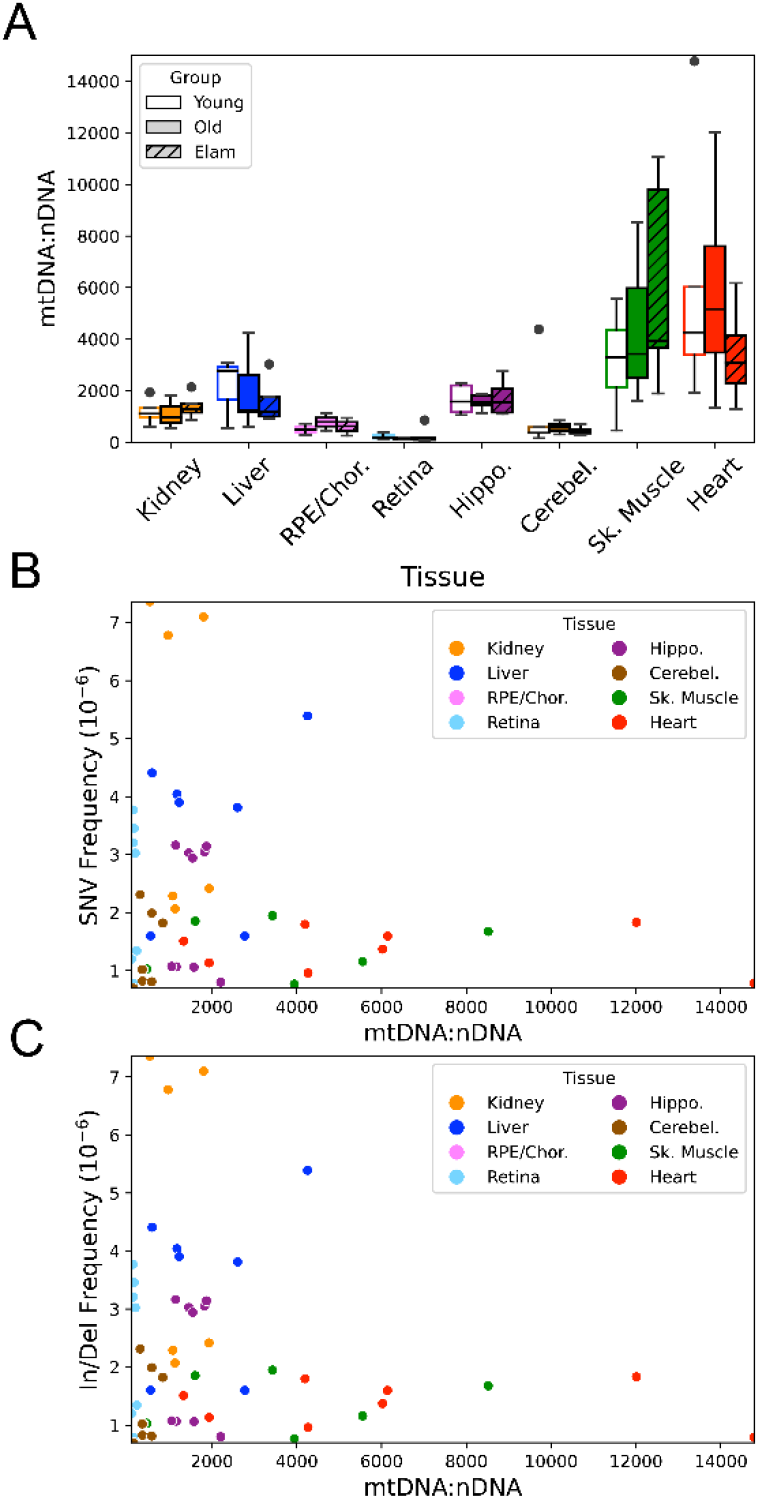
mtDNA copy shows no correlation with age, intervention, or mutation frequency. **(A)** mtDNA:nDNA ratio varies considerably between the eight tissue types, but does not change with age or treatment with Elam. **(B)** SNV mutation frequency and **(C)** In/Del mutation frequency do not correlate with mtDNA copy number, indicating that the tissue specific differences in mtDNA frequency is not due to mtDNA content.

**Figure S4.**
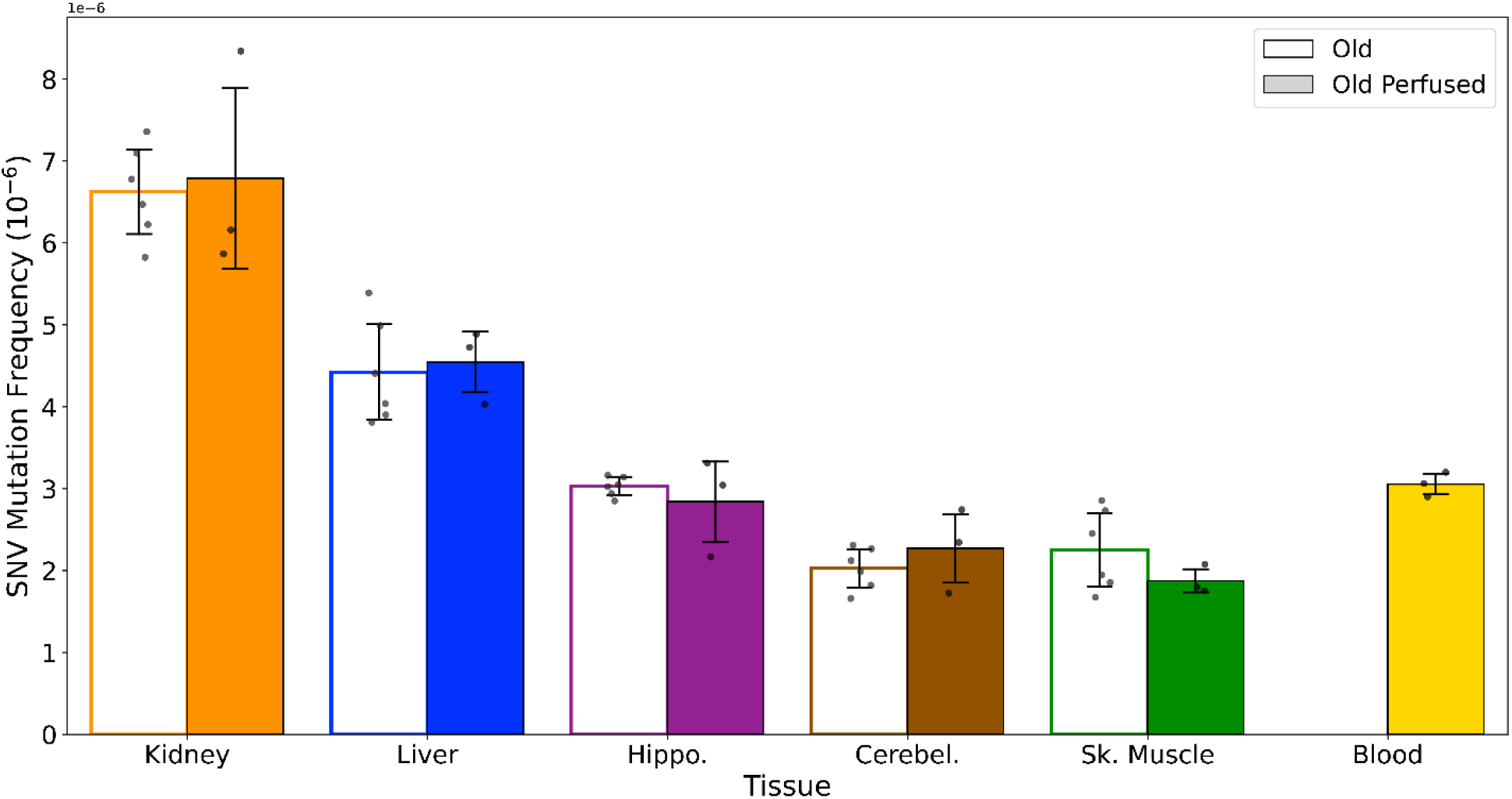
Blood does not significantly contribute to the differences in mutation frequency observed across tissues. A separate cohort of NIA male aged mice (26 mo, N=3) were transcardially perfused with 1x PBS prior to organ harvest. Collected tissues (kidney, liver, hippocampus, cerebellum, and skeletal muscle, and whole blood) were processed for Duplex-Seq using the same protocol as used in the main aging cohorts.

**Figure S5.**
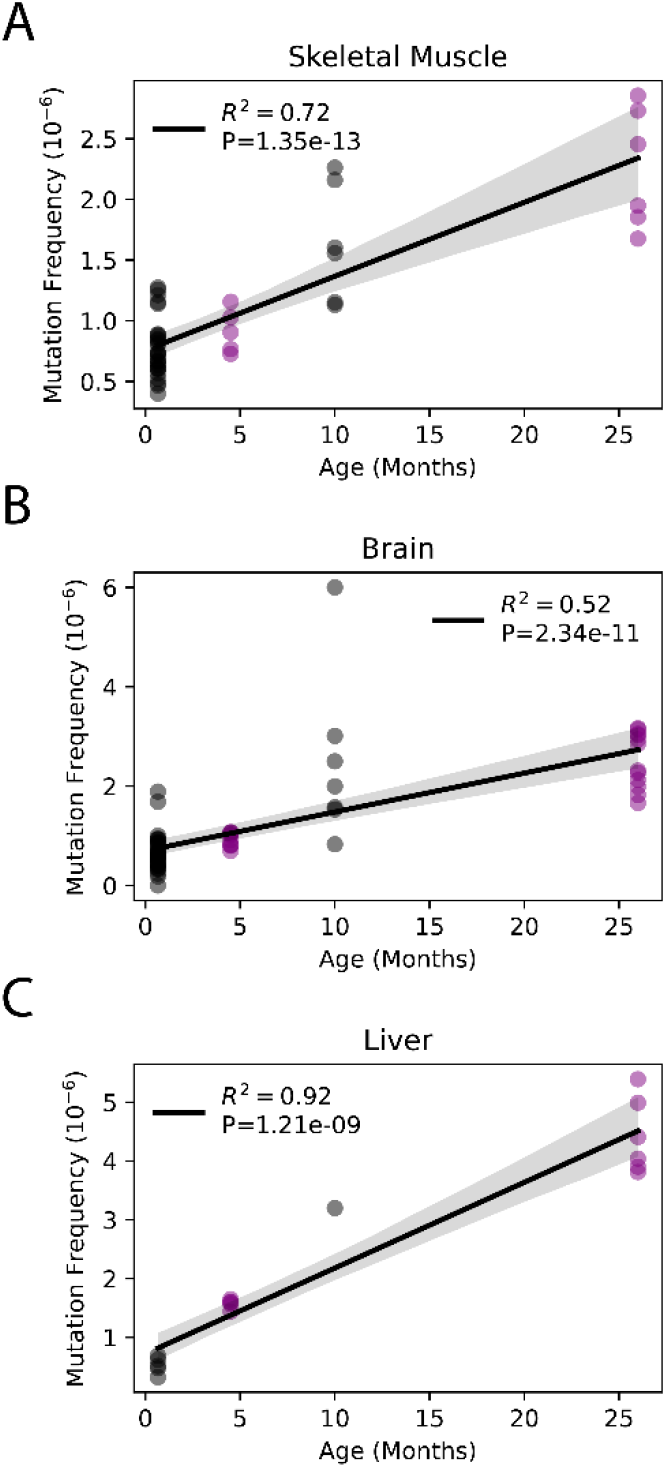
Somatic SNV mutations increase linearly with age. Linear regression of total SNV mutation frequency *vs* age in **(A)** skeletal muscle, **(B)** brain, and **(C)** liver. *black=*data from Arbeithuber et al.; *purple*=data from this study; *shaded area*=95% confidence interval of linear regression.

**Figure S6.**
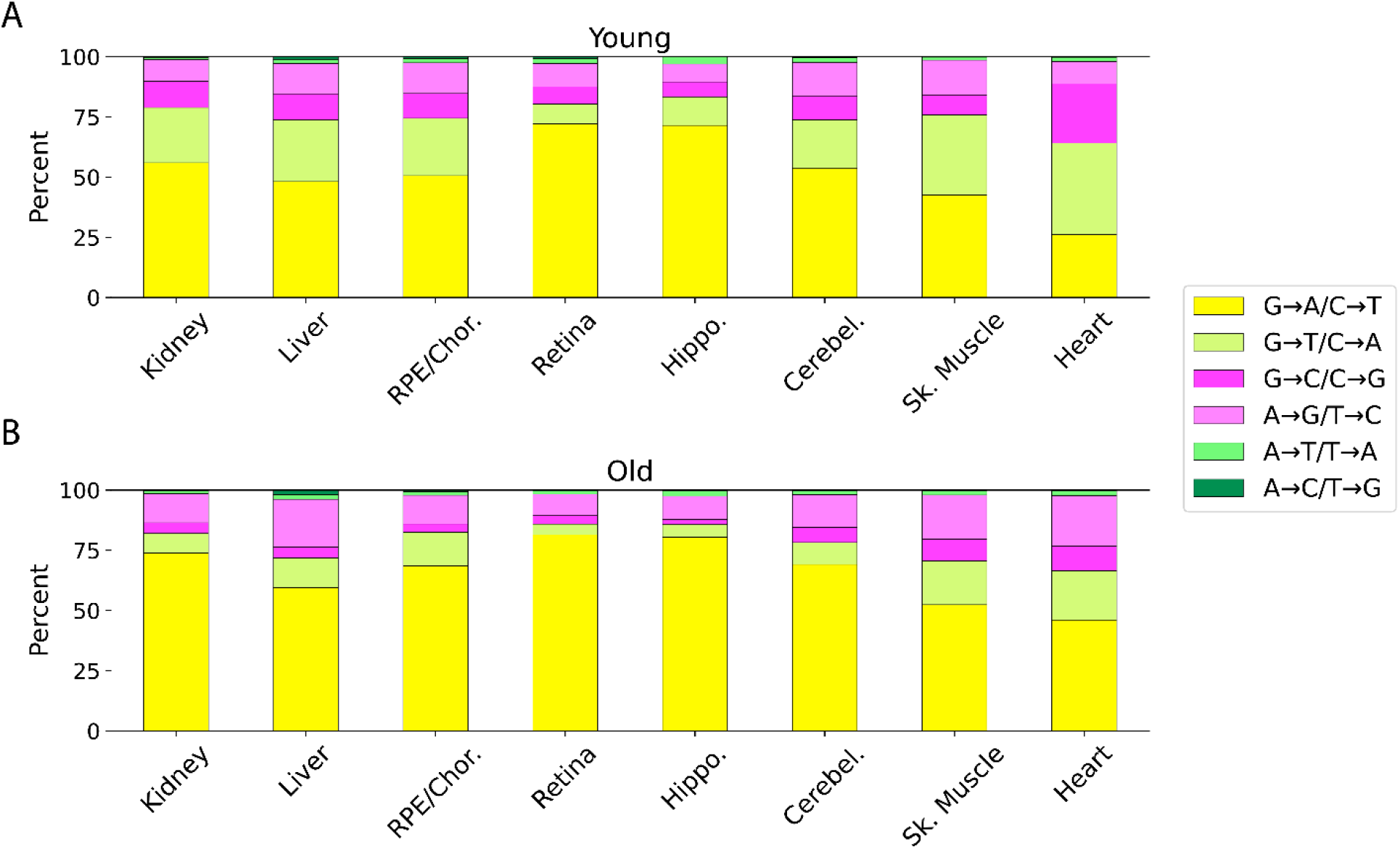
Relative proportion of different mutation types varies across tissues. **(A)** Young; **(B)** Old.

**Figure S7.**
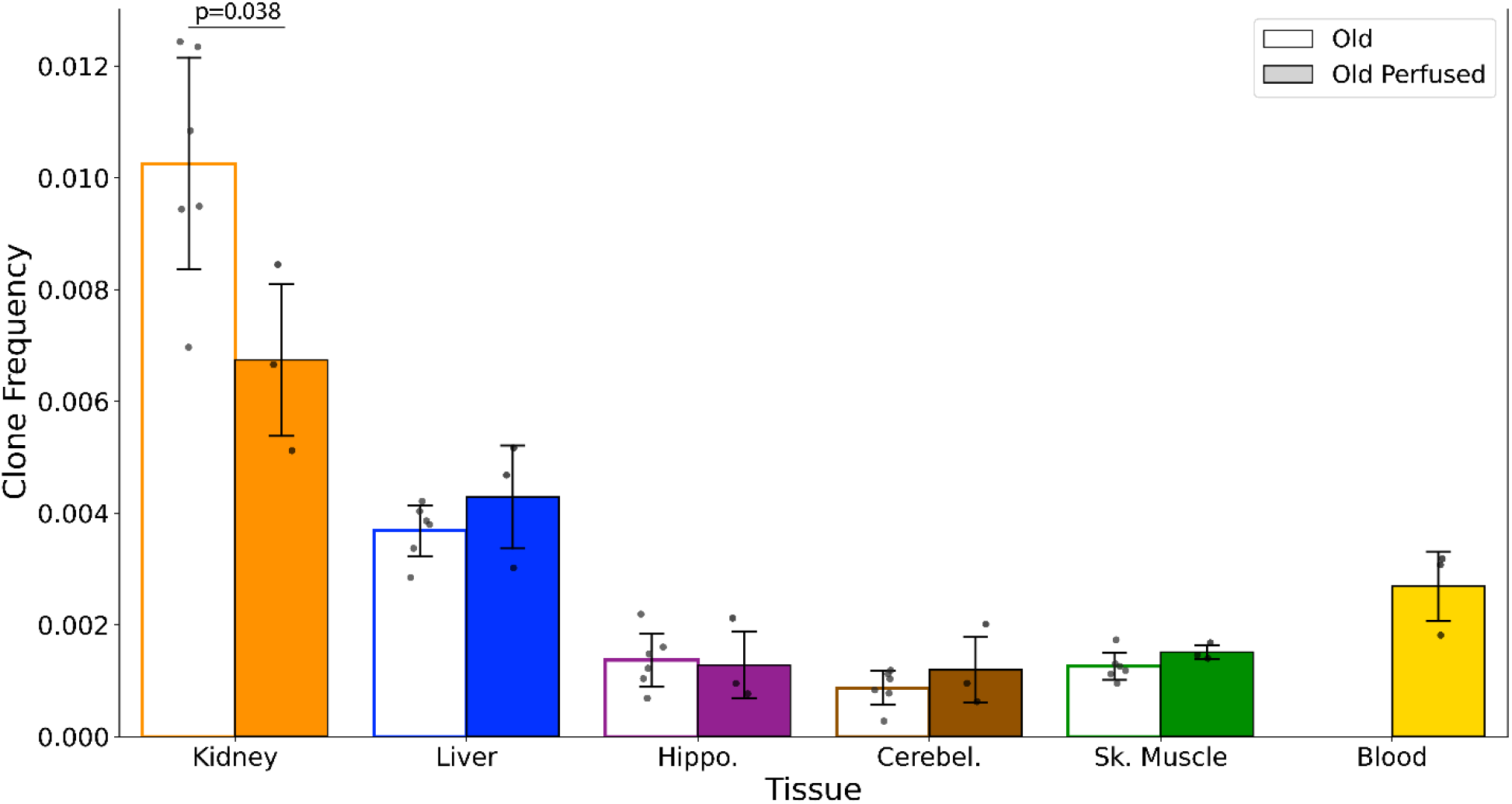
mtDNA variant clones in blood are not a significant contributor to age-related to clonal expansions. A separate cohort of NIA male aged mice (26 mo, N=3) were transcardially perfused with 1x PBS prior to organ harvest. Collected tissues (kidney, liver, hippocampus, cerebellum, and skeletal muscle, and whole blood) were processed for Duplex-Seq using the same protocol as used in the main aging cohorts.

**Figure S8.**
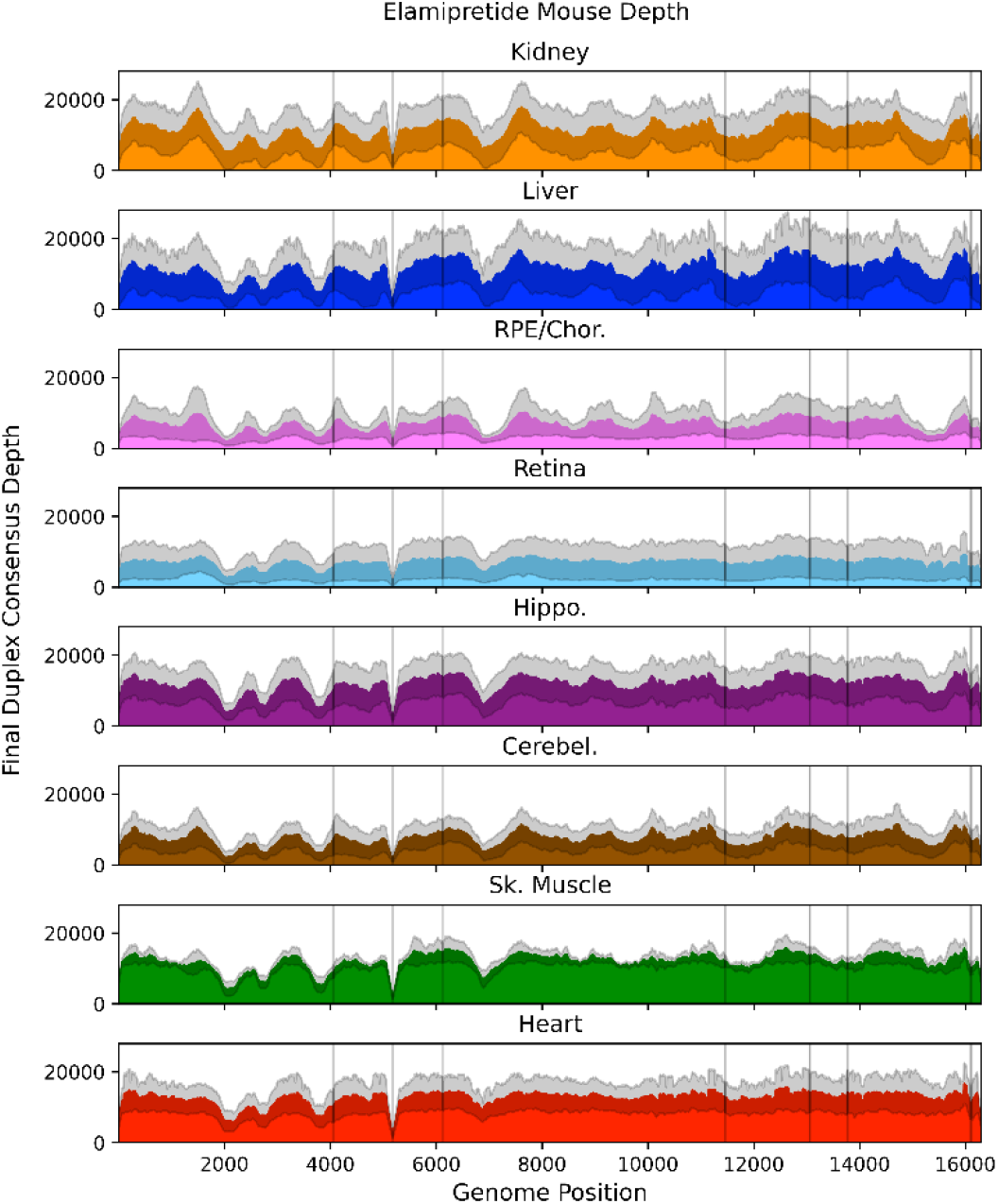
Mean post-consensus ‘Duplex’ depth for aged (26 month) Elamipretide treated tissues. Variant occurring within the masked regions (*vertical lines*) or positions with less than a post-consusensus depth of 100X were ignored. *Grey shading* = standard deviation of the mean for N=5 mice.

**Figure S9.**
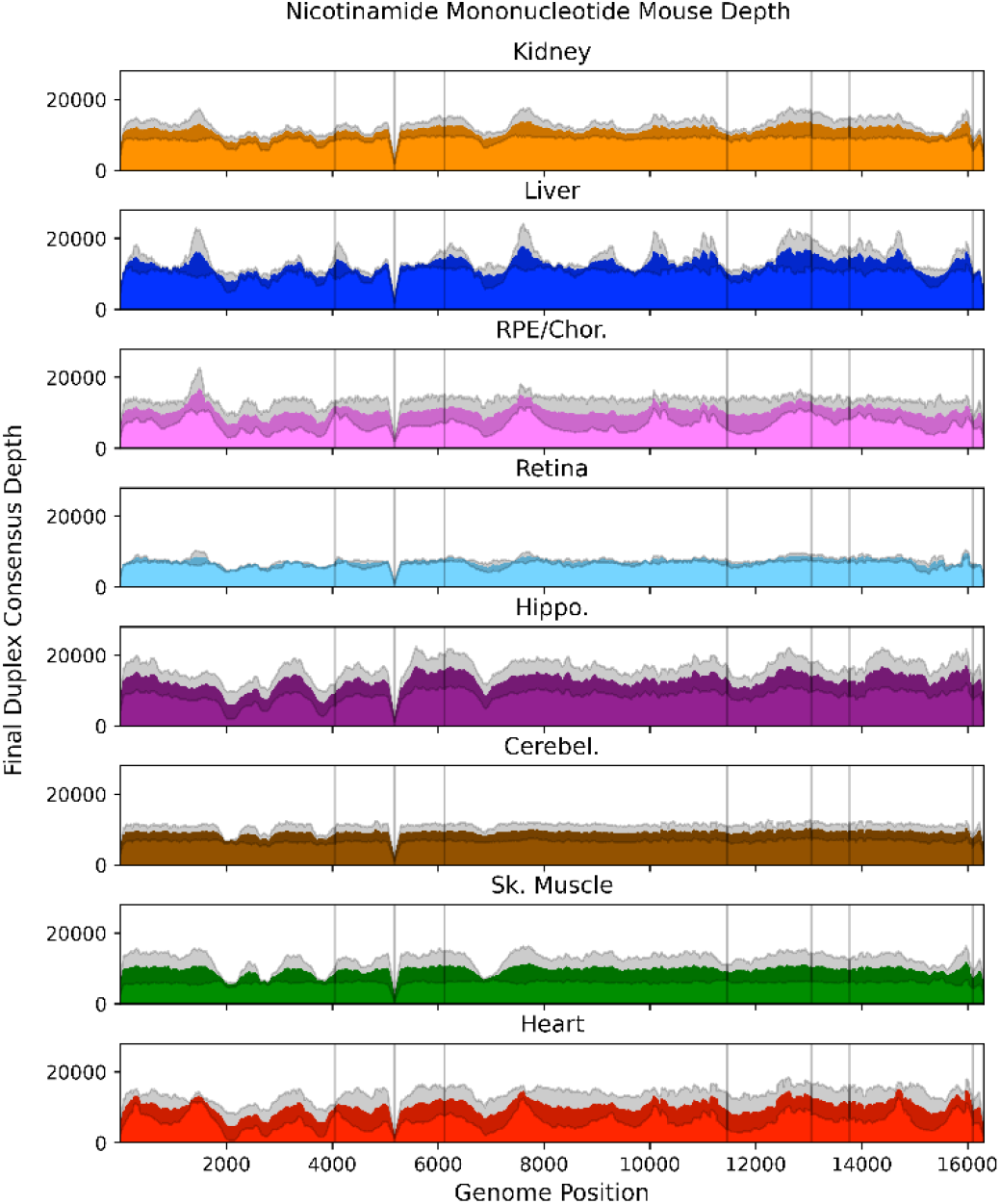
Mean post-consensus ‘Duplex’ depth for aged (26 month) nicotinamide mononucleotide treated tissues. Variant occurring within the masked regions (*vertical lines*) or positions with less than a post-consusensus depth of 100X were ignored. *Grey shading* = standard deviation of the mean for N=3 mice.

**Figure S10.**
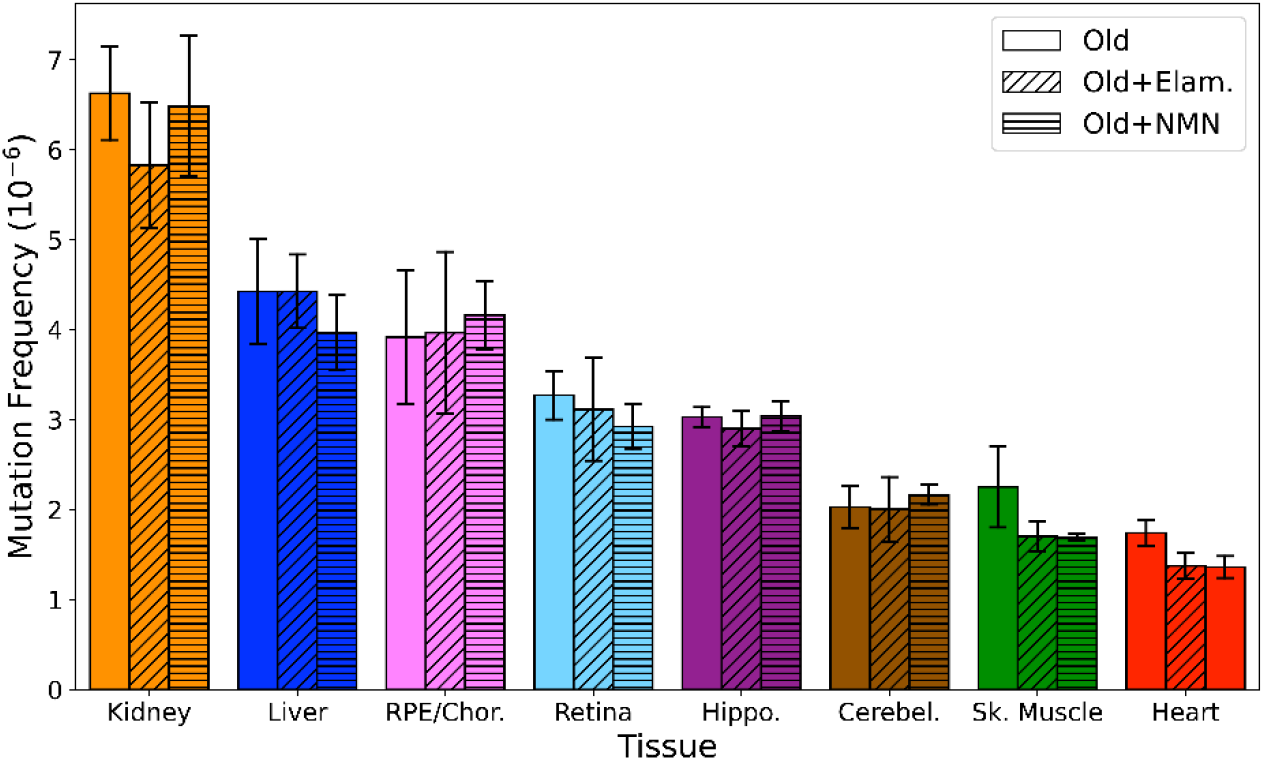
Elamipretide and nicotinamide mononucleotide do not affect the overall mtDNA mutation frequency. Overall point mutation frequency for old, old+Elam, and old+NMN separated by tissue. Dunnett’s test with untreated old as the control was used to test for significance. Error bars=standard deviation.

**Figure S11.**
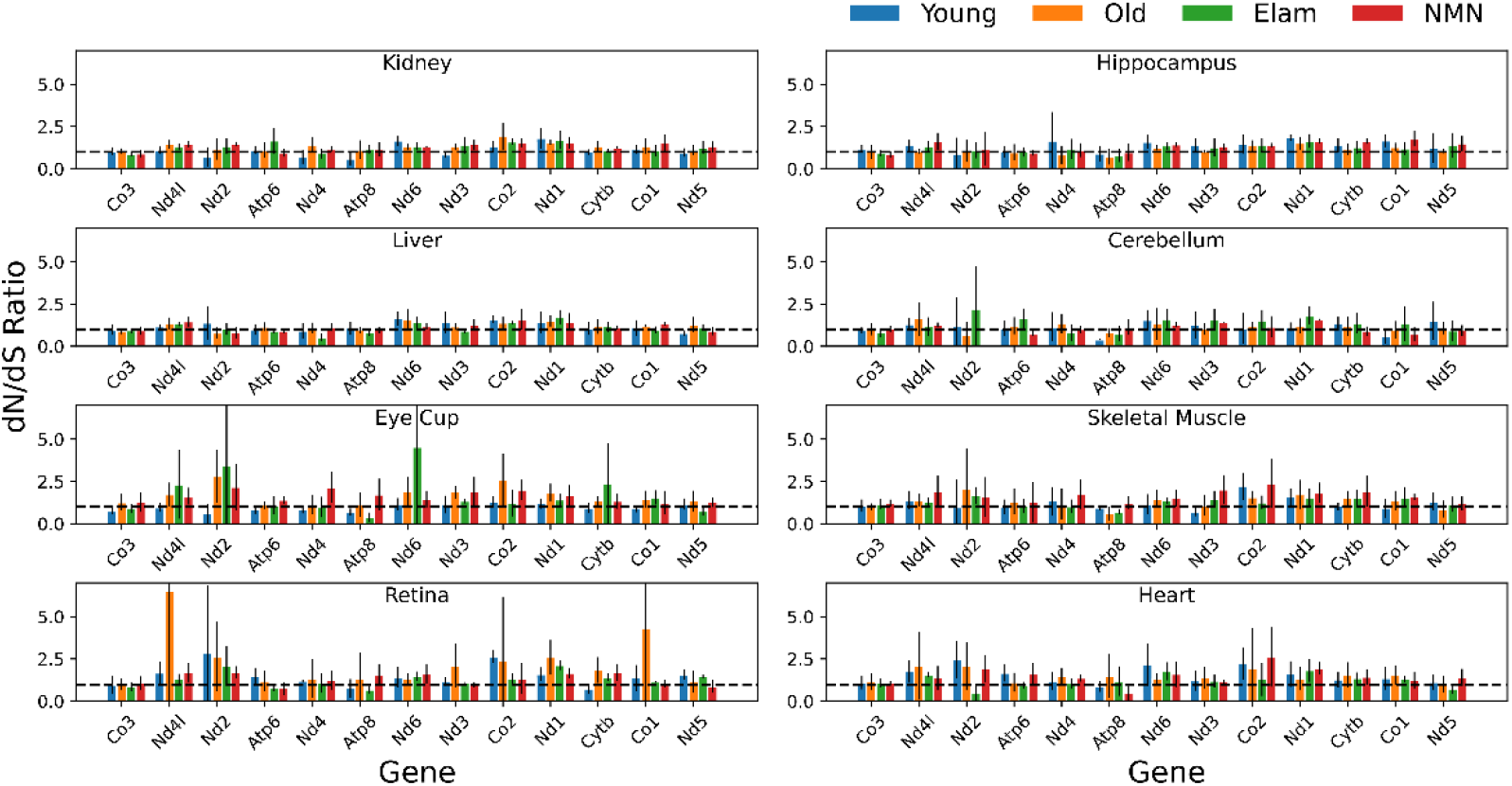
Per gene dN/dS ratio shows no apparent selection across age, tissues, and interventions. Variants for each sample were separated by protein coding gene and the dN/dS ratio calculated using the *dNdScv* R package using the median depth for each gene as a covariate. dN/dS ratios from the same tissue and age or treatment group were averaged. Error bars=standard deviation.

**Figure S12.**
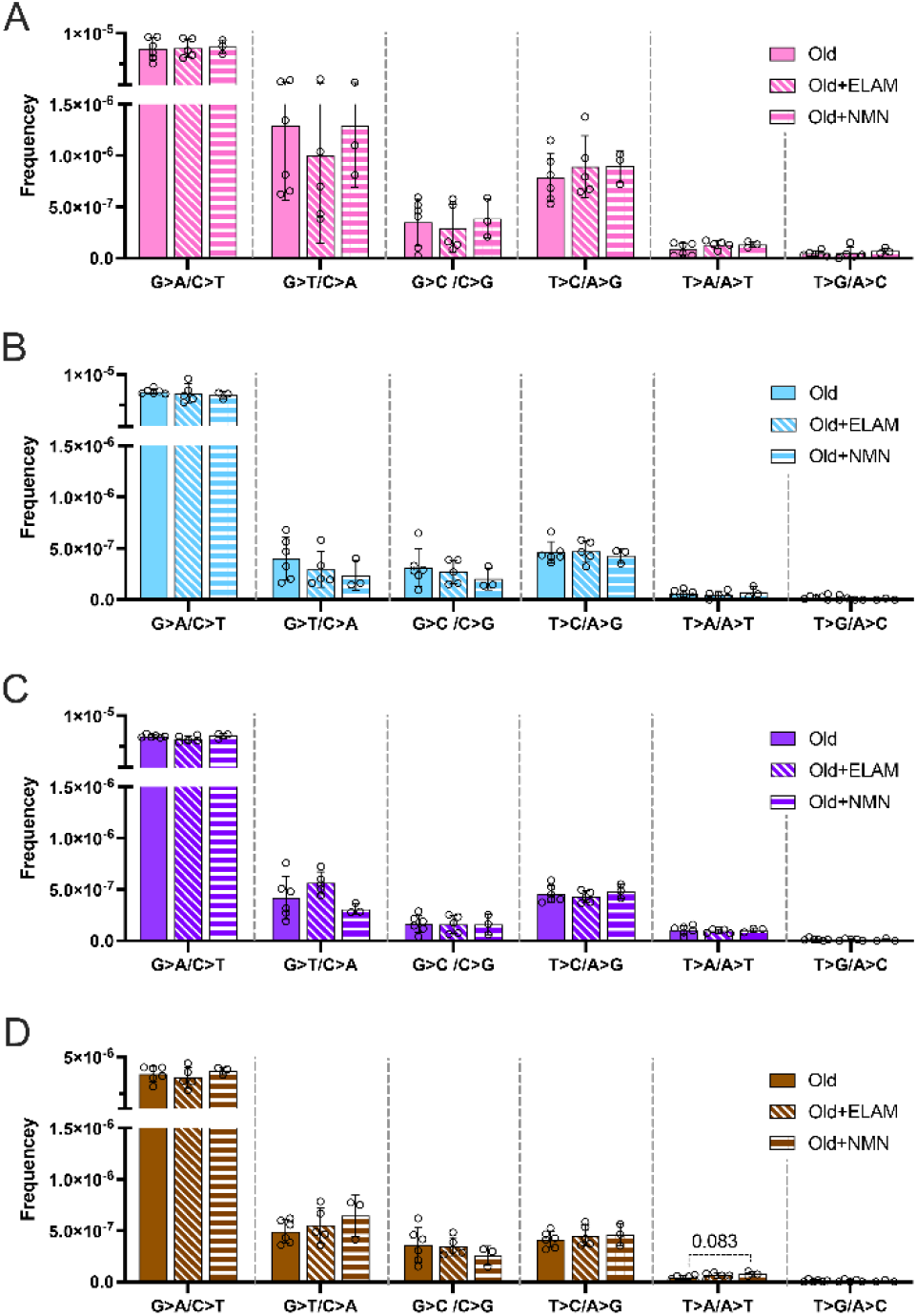
Late-life treatment with mitochondrially-targeted interventions does not reduce somatic mtDNA mutation burden in RPE/choroid (pink), Retina (cyan blue), Hippocampus (purple) or Cerebellum (brown). One-way ANOVA for each mutation class within tissue, and Dunnett’s multiple comparison test compared to untreated aged control group was performed with no significant differences or trends detected.

**Table S1. Sequence of the 96 defined UMI Duplex Sequencing adapters**. Sequence is provided in 5’→3’ orientation. Complementary UMI sequences are highlighted in red.

**Table S2. Summary of Duplex Sequencing data**. Summary of the samples sequenced, including assay performance metrics, including mtDNA enrichment specificity, family size, and consensus metrics, bases sequenced, sequencing depth, mutation counts, and mutation frequencies.

**Table S3. mtDNA to nDNA copy number ratio data**. Summary of the samples used to determine mtDNA and nDNA copy numbers. ND=not determined

**Table S4. Genome coordinates of regions masked in the analysis**. Coordinates are 1-indexed and columns are in .bed format order.

## References

Abascal F, Harvey LM, Mitchell E, Lawson ARJ, Lensing SV, Ellis P, Russell AJ, Alcantara RE, Baez-Ortega A, Wang Y, Kwa EJ, Lee-Six H, Cagan A, Coorens TH, Chapman MS, Olafsson S, Leonard S, Jones D, Machado HE, Davies M, Øbro NF, Mahubani KT, Allinson K, Gerstung M, Saeb-Parsy K, David G Kent, Elisa Laurenti, Stratton MR, Rahbari R, Campbell PJ, Osborne RJ, Martincorena I. 2021. Somatic mutation landscapes at single-molecule resolution. Nature 593:405–410.

Alexandrov LB, Jones PH, Wedge DC, Sale JE, Campbell PJ, Nik-Zainal S, Stratton MR. 2015. Clock-like mutational processes in human somatic cells. Nat Genet 47:1402–1407. doi:10.1038/ng.3441

Ameur A, Stewart JB, Freyer C, Hagström E, Ingman M, Larsson N-G, Gyllensten U. 2011. Ultra-deep sequencing of mouse mitochondrial DNA: Mutational patterns and their origins. PLoS Genet 7:e1002028. doi:10.1371/journal.pgen.1002028

Andreazza S, Samstag CL, Sanchez-Martinez A, Fernandez-Vizarra E, Gomez-Duran A, Lee JJ, Tufi R, Hipp MJ, Schmidt EK, Nicholls TJ, Gammage PA, Chinnery PF, Minczuk M, Pallanck LJ, Kennedy SR, Whitworth AJ. 2019. Mitochondrially-targeted APOBEC1 is a potent mtDNA mutator affecting mitochondrial function and organismal fitness in Drosophila. Nat Commun 10:3280. doi:10.1038/s41467-019-10857-y

Arbeithuber B, Hester J, Cremona MA, Stoler N, Zaidi A, Higgins B, Anthony K, Chiaromonte F, Diaz FJ, Makova KD. 2020. Age-related accumulation of de novo mitochondrial mutations in mammalian oocytes and somatic tissues. PLoS Biol 18:e3000745. doi:10.1371/journal.pbio.3000745

Campbell MD, Duan J, Samuelson AT, Gaffrey MJ, Merrihew GE, Egertson JD, Wang L, Bammler TK, Moore RJ, White CC, Kavanagh TJ, Voss JG, Szeto HH, Rabinovitch PS, MacCoss MJ, Qian W-J, Marcinek DJ. 2019. Improving mitochondrial function with SS-31 reverses age-related redox stress and improves exercise tolerance in aged mice. Free Radic Biol Med 134:268–281. doi:10.1016/j.freeradbiomed.2018.12.031

Chen H, Vermulst M, Wang YE, Chomyn A, Prolla TA, McCaffery JM, Chan DC. 2010. Mitochondrial fusion is required for mtDNA stability in skeletal muscle and tolerance of mtDNA mutations. Cell 141:280–289. doi:10.1016/j.cell.2010.02.026

Chen Z, Qi Y, French S, Zhang G, Garcia RC, Balaban R, Xu H. 2015. Genetic mosaic analysis of a deleterious mitochondrial DNA mutation in Drosophila reveals novel aspects of mitochondrial regulation and function. Mol Biol Cell 26:674–684. doi:10.1091/mbc.E14-11-1513

Chiao YA, Zhang H, Sweetwyne M, Whitson J, Ting YS, Basisty N, Pino LK, Quarles E, Nguyen N-H, Campbell MD, Zhang T, Gaffrey MJ, Merrihew G, Wang L, Yue Y, Duan D, Granzier HL, Szeto HH, Qian W-J, Marcinek D, MacCoss MJ, Rabinovitch P. 2020. Late-life restoration of mitochondrial function reverses cardiac dysfunction in old mice. eLife 9:e55513. doi:10.7554/eLife.55513

Colom B, Alcolea MP, Piedrafita G, Hall MWJ, Wabik A, Dentro SC, Fowler JC, Herms A, King C, Ong SH, Sood RK, Gerstung M, Martincorena I, Hall BA, Jones PH. 2020. Spatial competition shapes the dynamic mutational landscape of normal esophageal epithelium. Nat Genet 52:604–614. doi:10.1038/s41588-020-0624-3

Felsenstein J. 1974. The evolutionary advantage of recombination. Genetics 78:737–756. doi:10.1093/genetics/78.2.737

Fernández-Vizarra E, Enríquez JA, Pérez-Martos A, Montoya J, Fernández-Silva P. 2011. Tissue-specific differences in mitochondrial activity and biogenesis. Mitochondrion 11:207–213. doi:10.1016/j.mito.2010.09.011

Fieller E. 1954. Some problems in interval estimation. J Royal Stat Soc Ser B 16:175–185.

Gitschlag BL, Kirby CS, Samuels DC, Gangula RD, Mallal SA, Patel MR. 2016. Homeostatic responses regulate selfish mitochondrial genome dynamics in C. elegans. Cell Metab 24:91–103. doi:10.1016/j.cmet.2016.06.008

Greaves LC, Barron MJ, Plusa S, Kirkwood TB, Mathers JC, Taylor RW, Turnbull DM. 2010. Defects in multiple complexes of the respiratory chain are present in ageing human colonic crypts. Exp Gerontol 45:573–579. doi:10.1016/j.exger.2010.01.013

Greaves LC, Nooteboom M, Elson JL, Tuppen HAL, Taylor GA, Commane DM, Arasaradnam RP, Khrapko K, Taylor RW, Kirkwood TBL, Mathers JC, Turnbull DM. 2014. Clonal expansion of early to mid-life mitochondrial DNA point mutations drives mitochondrial dysfunction during human ageing. PLoS Genet 10:e1004620. doi:10.1371/journal.pgen.1004620

Greaves LC, Preston SL, Tadrous PJ, Taylor RW, Barron MJ, Oukrif D, Leedham SJ, Deheragoda M, Sasieni P, Novelli MR, Jankowski JAZ, Turnbull DM, Wright NA, McDonald SAC. 2006. Mitochondrial DNA mutations are established in human colonic stem cells, and mutated clones expand by crypt fission. Proc Natl Acad Sci USA 103:714–719. doi:10.1073/pnas.0505903103

Guan Y, Wang S-R, Huang X-Z, Xie Q, Xu Y-Y, Shang D, Hao C-M. 2017. Nicotinamide mononucleotide, an NAD+ precursor, rescues age-associated susceptibility to AKI in a sirtuin 1–dependent manner. J Am Soc Nephrol 28:2337–2352. doi:10.1681/ASN.2016040385

Hoekstra JG, Hipp MJ, Montine TJ, Kennedy SR. 2016. Mitochondrial DNA mutations increase in early stage Alzheimer disease and are inconsistent with oxidative damage. Ann Neurol 80:301–306. doi:10.1002/ana.24709

Horvath S. 2013. DNA methylation age of human tissues and cell types. Genome Biol 14:R115. doi:10.1186/gb-2013-14-10-r115

Itsara LS, Kennedy SR, Fox EJ, Yu S, Hewitt JJ, Sanchez-Contreras M, Cardozo-Pelaez F, Pallanck LJ. 2014. Oxidative stress is not a major contributor to somatic mitochondrial DNA mutations. PLoS Genet 10:e1003974. doi:10.1371/journal.pgen.1003974

Ju YS, Alexandrov LB, Gerstung M, Martincorena I, Nik-Zainal S, Ramakrishna M, Davies HR, Papaemmanuil E, Gundem G, Shlien A, Bolli N, Behjati S, Tarpey PS, Nangalia J, Massie CE, Butler AP, Teague JW, Vassiliou GS, Green AR, D. M-Q, Unnikrishnan A, Pimanda JE, Teh BT, Munshi N, Greaves M, Vyas P, El-Naggar AK, Santarius T, Collins VP, Grundy R, Taylor JA, Hayes DN, Malkin D, ICGC Breast Cancer Group, ICGC Chronic Myeloid Disorders Group, ICGC Prostate Cancer Group, Foster CS, Warren AY, Whitaker HC, Brewer D, Eeles R, Cooper C, Neal D, Visakorpi T, Isaacs WB, Bova GS, Flanagan AM, Futreal PA, Lynch AG, Chinnery PF, McDermott U, Stratton MR, Campbell PJ. 2014. Origins and functional consequences of somatic mitochondrial DNA mutations in human cancer. Elife 3:e02935. doi:10.7554/eLife.02935

Kauppila JHK, Bonekamp NA, Mourier A, Isokallio MA, Just A, Kauppila TES, Stewart JB, Larsson N-G. 2018. Base-excision repair deficiency alone or combined with increased oxidative stress does not increase mtDNA point mutations in mice. Nucleic Acids Res 46:6642– 6669. doi:10.1093/nar/gky456

Kauppila JHK, Stewart JB. 2015. Mitochondrial DNA: Radically free of free-radical driven mutations. Biochim Biophys Acta 1847:1354–1361. doi:10.1016/j.bbabio.2015.06.001

Kennedy SR, Salk JJ, Schmitt MW, Loeb LA. 2013. Ultra-sensitive sequencing reveals an age-related increase in somatic mitochondrial mutations that are inconsistent with oxidative damage. PLoS Genet 9:e1003794. doi:10.1371/journal.pgen.1003794

Kennedy SR, Schmitt MW, Fox EJ, Kohrn BF, Salk JJ, Ahn EH, Prindle MJ, Kuong KJ, Shen J-C, Risques R-A, Loeb LA. 2014. Detecting ultralow-frequency mutations by Duplex Sequencing. Nat Protoc 9:2586–2606. doi:10.1038/nprot.2014.170

Kowaltowski AJ. 2000. Alternative mitochondrial functions in cell physiopathology: beyond ATP production. Braz J Med Biol Res 33:241–250. doi:10.1590/S0100-879X2000000200014

Lareau CA, Ludwig LS, Sankaran VG. 2019. Longitudinal assessment of clonal mosaicism in human hematopoiesis via mitochondrial mutation tracking. Blood Adv 3:4161–4165. doi:10.1182/bloodadvances.2019001196

Larsson N-G. 2010. Somatic mitochondrial DNA mutations in mammalian aging. Annu Rev Biochem 79:683–706. doi:10.1146/annurev-biochem-060408-093701

Li R, Di L, Li J, Fan W, Liu Y, Guo W, Liu W, Liu L, Li Q, Chen L, Chen Y, Miao C, Liu H, Wang Y, Ma Y, Xu D, Lin D, Huang Y, Wang J, Bai F, Wu C. 2021. A body map of somatic mutagenesis in morphologically normal human tissues. Nature 597:398–403. doi:10.1038/s41586-021-03836-1

Lin Y-F, Schulz AM, Pellegrino MW, Lu Y, Shaham S, Haynes CM. 2016. Maintenance and propagation of a deleterious mitochondrial genome by the mitochondrial unfolded protein response. Nature 533:416–419. doi:10.1038/nature17989

López-Otín C, Blasco MA, Partridge L, Serrano M, Kroemer G. 2013. The hallmarks of aging. Cell 153:1194–1217. doi:10.1016/j.cell.2013.05.039

Ma H, Lee Y, Hayama T, Van Dyken C, Marti-Gutierrez N, Li Y, Ahmed R, Koski A, Kang E, Darby H, Gonmanee T, Park Y, Wolf DP, Jai Kim C, Mitalipov S. 2018. Germline and somatic mtDNA mutations in mouse aging. PLoS ONE 13:e0201304. doi:10.1371/journal.pone.0201304

Marcelino LA, Thilly WG. 1999. Mitochondrial mutagenesis in human cells and tissues. Mutat Res 434:177–203. doi:10.1016/S0921-8777(99)00028-2

Martin AS, Abraham DM, Hershberger KA, Bhatt DP, Mao L, Cui H, Liu J, Liu X, Muehlbauer MJ, Grimsrud PA, Locasale JW, Payne RM, Hirschey MD. 2017. Nicotinamide mononucleotide requires SIRT3 to improve cardiac function and bioenergetics in a Friedreich’s ataxia cardiomyopathy model. JCI Insight 2:e93885. doi:10.1172/jci.insight.93885

Martincorena I, Fowler JC, Wabik A, Lawson ARJ, Abascal F, Hall MWJ, Cagan A, Murai K, Mahbubani K, Stratton MR, Fitzgerald RC, Handford PA, Campbell PJ, Saeb-Parsy K, Jones PH. 2018. Somatic mutant clones colonize the human esophagus with age. Science 362:911–917. doi:10.1126/science.aau3879

Martincorena I, Raine KM, Gerstung M, Dawson KJ, Haase K, Van Loo P, Davies H, Stratton MR, Campbell PJ. 2017. Universal patterns of selection in cancer and somatic tissues. Cell 171:1029-1041.e21. doi:10.1016/j.cell.2017.09.042

Martincorena I, Roshan A, Gerstung M, Ellis P, Van Loo P, McLaren S, Wedge DC, Fullam A, Alexandrov LB, Tubio JM, Stebbings L, Menzies A, Widaa S, Stratton MR, Jones PH, Campbell PJ. 2015. High burden and pervasive positive selection of somatic mutations in normal human skin. Science 348:880–886. doi:10.1126/science.aaa6806

Masser DR, Clark NW, Van Remmen H, Freeman WM. 2016. Loss of the antioxidant enzyme CuZnSOD (Sod1) mimics an age-related increase in absolute mitochondrial DNA copy number in the skeletal muscle. Age (Dordr) 38:323–333. doi:10.1007/s11357-016-9930-1

Mitchell W, Ng EA, Tamucci JD, Boyd KJ, Sathappa M, Coscia A, Pan M, Han X, Eddy NA, May ER, Szeto HH, Alder NN. 2020. The mitochondria-targeted peptide SS-31 binds lipid bilayers and modulates surface electrostatics as a key component of its mechanism of action. J Biol Chem 295:7452–7469. doi:10.1074/jbc.RA119.012094

Muller H. 1964. The relation of recombination to mutational advance. Mutat Res 1:2–9. doi:10.1016/0027-5107(64)90047-8

Nekhaeva E, Bodyak ND, Kraytsberg Y, McGrath SB, Van Orsouw NJ, Pluzhnikov A, Wei JY, Vijg J, Khrapko K. 2002. Clonally expanded mtDNA point mutations are abundant in individual cells of human tissues. Proc Natl Acad Sci USA 99:5521–5526. doi:10.1073/pnas.072670199

Obi C, Smith AT, Hughes GJ, Adeboye AA. 2022. Targeting mitochondrial dysfunction with elamipretide. Heart Fail Rev. doi:10.1007/s10741-021-10199-2

Pickles S, Vigié P, Youle RJ. 2018. Mitophagy and quality control mechanisms in mitochondrial maintenance. Curr Biol 28:R170–R185. doi:10.1016/j.cub.2018.01.004

Pickrell AM, Huang C-H, Kennedy SR, Ordureau A, Sideris DP, Hoekstra JG, Harper JW, Youle RJ. 2015. Endogenous Parkin preserves dopaminergic substantia nigral neurons following mitochondrial DNA mutagenic stress. Neuron 87:371–381. doi:10.1016/j.neuron.2015.06.034

Rossignol R, Faustin B, Rocher C, Malgat M, Mazat J-P, Letellier T. 2003. Mitochondrial threshold effects. Biochem J 370:751–762. doi:10.1042/bj20021594

Rossignol R, Malgat M, Mazat J-P, Letellier T. 1999. Threshold effect and tissue specificity. J Biol Chem 274:33426–33432. doi:10.1074/jbc.274.47.33426

Salk JJ, Schmitt MW, Loeb LA. 2018. Enhancing the accuracy of next-generation sequencing for detecting rare and subclonal mutations. Nat Rev Genet 19:269–285. doi:10.1038/nrg.2017.117

Samstag CL, Hoekstra JG, Huang C-H, Chaisson MJ, Youle RJ, Kennedy SR, Pallanck LJ. 2018. Deleterious mitochondrial DNA point mutations are overrepresented in Drosophila expressing a proofreading-defective DNA polymerase γ. PLoS Genet 14:e1007805. doi:10.1371/journal.pgen.1007805

Samuels DC, Li C, Li B, Song Z, Torstenson E, Boyd Clay H, Rokas A, Thornton-Wells TA, Moore JH, Hughes TM, Hoffman RD, Haines JL, Murdock DG, Mortlock DP, Williams SM. 2013. Recurrent tissue-specific mtDNA mutations are common in humans. PLoS Genet 9:e1003929. doi:10.1371/journal.pgen.1003929

Sanchez-Contreras M, Kennedy SR. 2022. The complicated nature of somatic mtDNA mutations in aging. Front Aging 2:805126. doi:10.3389/fragi.2021.805126

Sanchez-Contreras M, Sweetwyne MT, Kohrn BF, Tsantilas KA, Hipp MJ, Schmidt EK, Fredrickson J, Whitson JA, Campbell MD, Rabinovitch PS, Marcinek DJ, Kennedy SR. 2021. A replication-linked mutational gradient drives somatic mutation accumulation and influences germline polymorphisms and genome composition in mitochondrial DNA. Nucl Acids Res 49:11103–11118.

Schmitt MW, Kennedy SR, Salk JJ, Fox EJ, Hiatt JB, Loeb LA. 2012. Detection of ultra-rare mutations by next-generation sequencing. Proc Natl Acad Sci USA 109:14508–14513. doi:10.1073/pnas.1208715109

Suen D-F, Narendra DP, Tanaka A, Manfredi G, Youle RJ. 2010. Parkin overexpression selects against a deleterious mtDNA mutation in heteroplasmic cybrid cells. Proc Natl Acad Sci USA 107:11835–11840. doi:10.1073/pnas.0914569107

Sweetwyne MT, Pippin JW, Eng DG, Hudkins KL, Chiao YA, Campbell MD, Marcinek DJ, Alpers CE, Szeto HH, Rabinovitch PS, Shankland SJ. 2017. The mitochondrial-targeted peptide, SS-31, improves glomerular architecture in mice of advanced age. Kidney Int 91:1126– 1145. doi:10.1016/j.kint.2016.10.036

Szczepanowska K, Trifunovic A. 2017. Origins of mtDNA mutations in ageing. Essays Biochem 61:325–337. doi:10.1042/EBC20160090

Vermulst M, Bielas JH, Kujoth GC, Ladiges WC, Rabinovitch PS, Prolla TA, Loeb LA. 2007.Mitochondrial point mutations do not limit the natural lifespan of mice. Nat Genet 39:540–543. doi:10.1038/ng1988

Wallace DC. 1999. Mitochondrial diseases in man and mouse. Science 283:1482–1488. doi:10.1126/science.283.5407.1482

Wei W, Tuna S, Keogh MJ, Smith KR, Aitman TJ, Beales PL, Bennett DL, Gale DP, Bitner-Glindzicz MAK, Black GC, Brennan P, Elliott P, Flinter FA, Floto RA, Houlden H, Irving M, Koziell A, Maher ER, Markus HS, Morrell NW, Newman WG, Roberts I, Sayer JA, Smith KGC, Taylor JC, Watkins H, Webster AR, Wilkie AOM, Williamson C, NIHR BioResource–Rare Diseases, 100,000 Genomes Project–Rare Diseases Pilot, Ashford S, Penkett CJ, Stirrups KE, Rendon A, Ouwehand WH, Bradley JR, Raymond FL, Caulfield M, Turro E, Chinnery PF. 2019. Germline selection shapes human mitochondrial DNA diversity. Science 364:eaau6520. doi:10.1126/science.aau6520

Whitson JA, Bitto A, Zhang H, Sweetwyne MT, Coig R, Bhayana S, Shankland EG, Wang L, Bammler TK, Mills KF, Imai S, Conley KE, Marcinek DJ, Rabinovitch PS. 2020. SS-31 and NMN: Two paths to improve metabolism and function in aged hearts. Aging Cell 19:e13213. doi:10.1111/acel.13213

Williams SL, Mash DC, Züchner S, Moraes CT. 2013. Somatic mtDNA mutation spectra in the aging human putamen. PLoS Genet 9:e1003990. doi:10.1371/journal.pgen.1003990

Yoshino J, Baur JA, Imai S. 2018. NAD+ intermediates: The biology and therapeutic potential of NMN and NR. Cell Metab 27:513–528. doi:10.1016/j.cmet.2017.11.002

Yoshino J, Mills KF, Yoon MJ, Imai S. 2011. Nicotinamide mononucleotide, a key NAD+ intermediate, treats the pathophysiology of diet- and age-induced diabetes in mice. Cell Metab 14:528–536. doi:10.1016/j.cmet.2011.08.014

Youle RJ, Narendra DP. 2011. Mechanisms of mitophagy. Nat Rev Mol Cell Biol 12:9–14. doi:10.1038/nrm3028

Zhang H, Alder NN, Wang W, Szeto H, Marcinek DJ, Rabinovitch PS. 2020. Reduction of elevated proton leak rejuvenates mitochondria in the aged cardiomyocyte. eLife 9:e60827. doi:10.7554/eLife.60827

Zhang L, Vijg J. 2018. Somatic mutagenesis in mammals and its implications for human disease and aging. Annu Rev Genet 52:397–419. doi:10.1146/annurev-genet-120417-031501

Zheng W, Khrapko K, Coller HA, Thilly WG, Copeland WC. 2006. Origins of human mitochondrial point mutations as DNA polymerase γ-mediated errors. Mutat Res 599:11–20. doi:10.1016/j.mrfmmm.2005.12.012

Zhu M, Lu T, Jia Y, Luo X, Gopal P, Li L, Odewole M, Renteria V, Singal AG, Jang Y, Ge K, Wang SC, Sorouri M, Parekh JR, MacConmara MP, Yopp AC, Wang T, Zhu H. 2019. Somatic mutations increase hepatic clonal fitness and regeneration in chronic liver disease. Cell 177:608-621.e12. doi:10.1016/j.cell.2019.03.026

